# mRNA detection in budding yeast with single fluorophores

**DOI:** 10.1101/155143

**Authors:** Gable M. Wadsworth, Rasesh Y. Parikh, John S. Choy, Harold D. Kim

## Abstract

Quantitative measurement of mRNA levels in single cells is necessary to understand phenotypic variability within an otherwise isogenic population of cells. Single-molecule mRNA Fluorescence In Situ Hybridization (FISH) has been established as the standard method for this purpose, but current protocols require a long region of mRNA to be targeted by multiple DNA probes. Here, we introduce a new single-probe FISH protocol termed sFISH for budding yeast, *Saccharomyces cerevisiae* using a single DNA probe labeled with a single fluorophore. In sFISH, we markedly improved probe specificity and signal-to-background ratio by using methanol fixation and inclined laser illumination. We show that sFISH reports mRNA changes that correspond to protein levels and gene copy number. Using this new FISH protocol, we can detect more than 50% of the total target mRNA. We also demonstrate the versatility of sFISH using FRET detection and mRNA isoform profiling as examples. Our FISH protocol with single-fluorophore sensitivity significantly reduces cost and time compared to the conventional FISH protocols and opens up new opportunities to investigate small changes in RNA at the single cell level.

## I. INTRODUCTION

Genetically identical cells are known to display phenotypic heterogeneity(1). This is due in large part to variation in gene expression from cell to cell(2, 3). Such single-cell variability can increase fitness of both unicellular and multicellular organisms against unpredictable changes in the environment(4) and may lead to other mechanisms of biological importance(5, 6). Hence, quantification of gene expression at the single cell level has become an active area of technological development. Transcription, which takes place in bursts at one or a few copies of a gene, is a significant source of gene expression noise, and therefore, measuring mRNA levels in single cells is essential to investigate noise generation mechanisms.

There are two general methods to quantify mRNA levels in single cells. The first type, which includes quantitative PCR (qPCR) and RNA sequencing (RNA-seq), relies on an indirect readout of DNA that is generated from reverse transcribed mRNA extracted from lysed cells. The main advantage of these methods is the scalability. Combined with a microfluidic or a single-cell sorting platform, these methods allow the whole transcriptome analysis of a single cell(7). However, these methods require laborious sample preparation and high-cost instruments, and are error-prone for short, low copy number transcripts(8). Due to the high cost of resources, these methods are in general not appropriate for a focused analysis of individual transcripts. These techniques also do not preserve information on subcellular localization of transcripts with respect to other cellular compartments, which may be valuable to understand their role and function(9).

In contrast, mRNA detection by fluorescence in situ hybridization (FISH) relies on a direct readout emanating from fluorescently labeled DNA probes bound to target mRNA molecules inside the cell. This method involves only low-cost cell processing and routine fluorescence microscopy and can yield absolute numbers for targeted transcripts without separate calibration steps. Since its first demonstration in 1982(10), mRNA FISH has undergone significant advancement, now capable of detecting single-nucleotide variants(11) and gene fusions(12, 13), and can be used to determine RNA sequence(14).

Currently, there are two widely used mRNA FISH protocols. The first protocol developed by the Singer lab(15, 16) uses ~ 50 nucleotide (nt) long probes that are each labeled with multiple fluorescent dyes. Five different probes, when hybridized to a single mRNA, generate an intense spot under a fluorescence microscope. The second protocol was introduced by Raj et al.(17), and features relatively short (~ 20 nt) probes that are singly labeled. To compensate for the smaller number of fluorophores per probe, 20-50 probes are used to detect a single mRNA. Both protocols require the target transcript to be quite long (> 200 nt) for hybridization of multiple probes(18, 19).

The need of multiple probes is two-fold. First, multiple probes increase the intensity of FISH spots. If *N* different probes can all bind to the same mRNA with equal probability, *p*, the mean number of probes per mRNA is *N_p_*. Secondly, using multiple probes increases the detection rate of mRNA because the probability of all *N* probes failing to bind an mRNA molecule is (1 − *p*)*^N^*. Therefore, in principle, one can decrease the false negatives (the number of undetected mRNA molecules) by increasing the number of probes.

However, the multiple probe requirement is not absolute. In many studies, single mRNA molecules were successfully probed with a smaller number of fluorophores: as few as four fluorophores for budding yeast(20) and a single fluorophore for E. coli(21). Recently, single nucleotide variants were detected at the single fluorophore level in human cells, albeit with multiple fluorophores of a different color used as a guide(11). Therefore, although considered difficult(22), detecting mRNA with a single fluorophore is achievable.

In many cases, mRNA FISH can be used for relative quantification of a specific transcript under different conditions or transcriptional heterogeneity in an isogenic population of cells. For such applications, the use of multiple probes is not needed. Reducing the number of probes may compromise the signal, but most FISH studies employ an epifluorescence microscope, which is not optimal for single-fluorophore detection with cellular background. We were thus motivated to revisit the current FISH method originally developed for budding yeast and explore the possibility of using single probes to measure relative abundance of mRNAs.

Here, we present single-fluorophore, single-probe FISH (sFISH) for budding yeast. In our modified method, a 26-nt probe with a single Cy3 or Cy5 dye is sufficient to accurately detect single mRNA molecules in single yeast cells. We employ methanol fixation and highly inclined laser illumination to increase the signal to noise ratio needed to capture fluorescence from single probes. We varied the transcription rate of a fluorescent reporter gene by changing the promoter strength and found that our new method could detect a proportional change in the corresponding mRNAs. By using three independent methods, we estimate the detection efficiency of this protocol at ~60 %. We anticipate that this method will add a highly useful tool for measuring steady-state transcript statistics of budding yeast. Technical improvements introduced in this protocol can also be used to study short transcripts in budding yeast.

## II. MATERIALS AND METHODS

### A. Strain construction

We constructed multiple PHO5 promoter variants of budding yeast following the protocol used in our previous study(23). These variants share a high affinity Pho4 binding site in the exposed region between nucleosome -2 and nucleosome -3, thus belonging to a family of HX promoter variants(24). These variants differ in their DNA sequence in the nucleosome -2 region (Supplementary Figure S1 and Supplementary Table S1). The PHO5 open reading frame was then replaced with a yellow fluorescent protein (yEVenus) gene by homologous recombination. To achieve constitutive expression of yEVenus, the PHO5 promoter variants were mated with the pho80Δ strain(25). yEVenus levels of these promoter variants were quantified using the epi-fluorescence microscope (Supplementary Figure S1B). Yeast strains with different ploidies (2n, 3n, 4n) were from Dr. David Pellman.

### B. Sample preparation

Our procedure closely follows the protocols by Youk et al.(26) and Raj et al. (27) with some modifications. Yeast cells are grown overnight to a final OD600 of 0.5 in 50mL of SD complete medium. Cells are fixed and then kept at 4°C. Fixation is performed either by treating the cells with 2 % v/v formaldehyde or with methanol for 10 minutes. In the case of methanol, cells are spun down, and the pellet is resuspended in methanol. The cells are washed twice with Buffer B (1.2 M sorbitol, 0.1 M potassium phosphate) at 4°C after fixation. Cells are then resuspended in 1mL of spheroplasting buffer (10mL Buffer B, 100 μL 200 mM vanadyl ribonucleoside complex from New England Biolabs). 2 μL of zymolyase at 5units/μL (Zymolyase-20T at 21000units/g from Seikagaku Business Corporation) are added into the mixture of cells, and then they are incubated at 30°C. The amount of digestion is determined by measuring OD600 of 1mL of a suspension containing 100 μL of the cell sample until the OD600 has decreased by about 30 %, which is typically 15 minutes. Cell wall digestion can be verified by allowing 10 μL cells to settle to the surface of a multi-well plate and adding 100 μL DI water to the well. Within a few minutes the majority of the cells should be lysed when inspected under a microscope. If the cells do not lyse, then the procedure has not effectively removed the cell wall. Following this treatment the cells are washed twice in Buffer B at 4 °C and stored in 70 % ethanol for at least one hour.

Hybridization is performed by washing the cells twice with 1mL of wash buffer containing 10 % formamide and 330 mM salt as SSC buffer. This step loosens the protein-RNA interactions to increase RNA accessibility for probe hybridization, and decreases nonspecific probe binding(28). Hybridization buffer is prepared in 10mL volumes containing 1mL 20X SSC (Ambion), 1mL formamide (Ambion), 100 μL 200 mM vanadyl ribonucleoside complex (New England Biolabs), 1 g dextran sulfate sodium salt (Sigma, D8906), 10 mg *Escherichia coli* tRNA (Sigma, R1753), 40 μL of 5 mg/mL BSA (Ambion), and 8mL deionized water. Cells are resuspended in hybridization buffer and the appropriate amount of probes is added to bring the working concentration to 65 nM and the final volume to 100 μL. Cells are then incubated at 30°C overnight. The following morning, cells are washed twice in wash buffer and left as a pellet. Slides are prepared by mixing 2 μL of concentrated cells with 2 μL of imaging buffer that has the same concentration of salt as the wash and hybridization buffers. 1.5 to 2 μL of cell suspension is placed between a microscope slide (1″ × 3″) and a coverslip (#1.5, 18 mm × 18 mm), and the chamber is gently pressed to form a monolayer of cells. The edges of the chamber are sealed with fast curing epoxy. The height of the chamber is estimated to be about 3 μm. Cells slightly squeezed in the chamber remain confined during the observation period. The imaging buffer contains the PCA/PCD oxygen scavenging system(29) to extend the photobleaching lifetime of the fluorophores. The imaging buffer is composed of 10 mM Tris at pH 8, 5 μL 20X SSC, 2.5 mM protocatechuic acid (PCA), 10 nM protocatechuate-3,4-dioxygenase (PCD), and 1 mM Trolox (6-hydroxy-2,5,7,8-tetramethylchroman-2-carboxylic acid).

### C. Microscope setup

Our custom setup was built around a commercial microscope (IX81, Olympus) equipped with motorized translation in z direction and a motorized filter wheel. We used 514 nm line from an Argon laser to excite yEVenus, a 532-nm solid state laser (LCX-532L-100, Oxxius) to excite Cy3, and a 640 nm solid state laser (1185055, Coherent) to excite Cy5. All lasers were coupled to a fiber optic, and the laser output from the fiber optic was collimated and focused at the back focal plane of the objective (Olympus UPlanSApo 100X/1.4 Oil). We mounted the fiber coupling onto a translational stage to vary the incidence angle of the excitation beam (Supplementary Figure S2).

### D. Imaging

The laser power was set to produce 25mW of power at the back focal plane of the objective. Z-stack images were acquired using an EMCCD (iXon plus, Andor) at 512x512 resolution every 0.2 μm step over 10 μm distance. The exposure time for the Cy5 channel was set to 100 ms. Hardware control and image acquisition were performed using Micromanager(30).

### E. Analysis

We used the MATLAB Image Processing Toolbox to batch process each three dimensional image through a series of automated steps(Supplementary Figure S3). Cell segmentation is accomplished by applying edge detection to DIC images of the cells. The detected edges are connected using [1x4] structural elements and binary morphological dilation and erosion operations. Cells are further selected based on the ellipticity and area of the detected regions.

Fluorescence intensities in cellular regions were background-corrected by subtracting the mean intensity taken from the cell boundary. Protein fluorescence was determined by taking the sum of the pixels inside the cell boundary and normalizing by the area. In FISH images, all maximum intensity pixels in the 3 × 3 × 3 neighborhood are selected as candidate spots. Spot intensity was calculated by taking the average of pixel values in a diffraction-limited ellipsoid and subtracting the background value evaluated from the pixels bounding the ellipsoid. The histogram of spot intensities was fit with a sum of three Gaussian distributions (Supplementary Figure S12). The first peak centered around zero arises mostly from noise. The second peak corresponds to spatially-resolved single transcripts. The third peak is likely due to multiple overlapping transcripts because this peak grows with gene expression level.

The important aspect of our spot identification algorithm is that spots are called using a locally determined threshold over a single cell. Due to nonuniform illumination, cells near the edge of the field of view are weakly excited, and the FISH spots are significantly dimmer than those near the center of the field of view. Therefore, applying cell-specific threshold can mitigate this problem, and reduce false negatives. To correct for the nonuniform illumination, we also adopted a post-acquisition algorithm called Corrected Intensity Distributions using Regularized Energy minimization (CIDRE)(31) prior to segmentation.

## III. RESULTS

### A. Observation of single fluorophores in vivo

While the current FISH approaches have been successfully applied to quantify mRNA level at the single cell level, they are not applicable to studies that investigate short RNA molecules or changes in short regions of RNA because of the multiple-probe requirement. To circumvent these limitations, we sought to develop a single-probe FISH technique for budding yeast. The three parameters we considered as we began our modified FISH method was the dye molecule, the fixative, and imaging technique. We chose Cy5 over Cy3 because yeast autofluorescence is lower in the red channel than the green(32). In addition, we tried two different fixatives that may differentially contribute to the signal-to-noise of Cy5(33).

We first acquired fluorescence images of cells of a positive control strain under epi-illumination. We were able to observe isolated fluorescent spots, but the signal-to-background ratio was poor. Hence, we tried illuminating the cells using an inclined geometry (Figure 1B). Highly inclined illumination excites a thin slice inside the cell, which leads to higher excitation intensity and lower out-of-focal plane background compared to epi-illumination(34). Using this setup, we observed a significant enhancement in the signal-to-noise ratio (Figure 1C-D). The oxygen scavenging system was also critical for detection of the isolated fluorescent spots. Two observations indicate that most of these spots arise from a single Cy5 molecule. First, their fluorescence intensities are comparable to the fluorescence intensity of single Cy5 molecules nonspecifically bound to the surface. Second, upon continuous excitation, most spots disappear in single photobleaching steps, consistent with a single Cy5 molecule (Supplementary Figure S5).

**FIG. 1.**
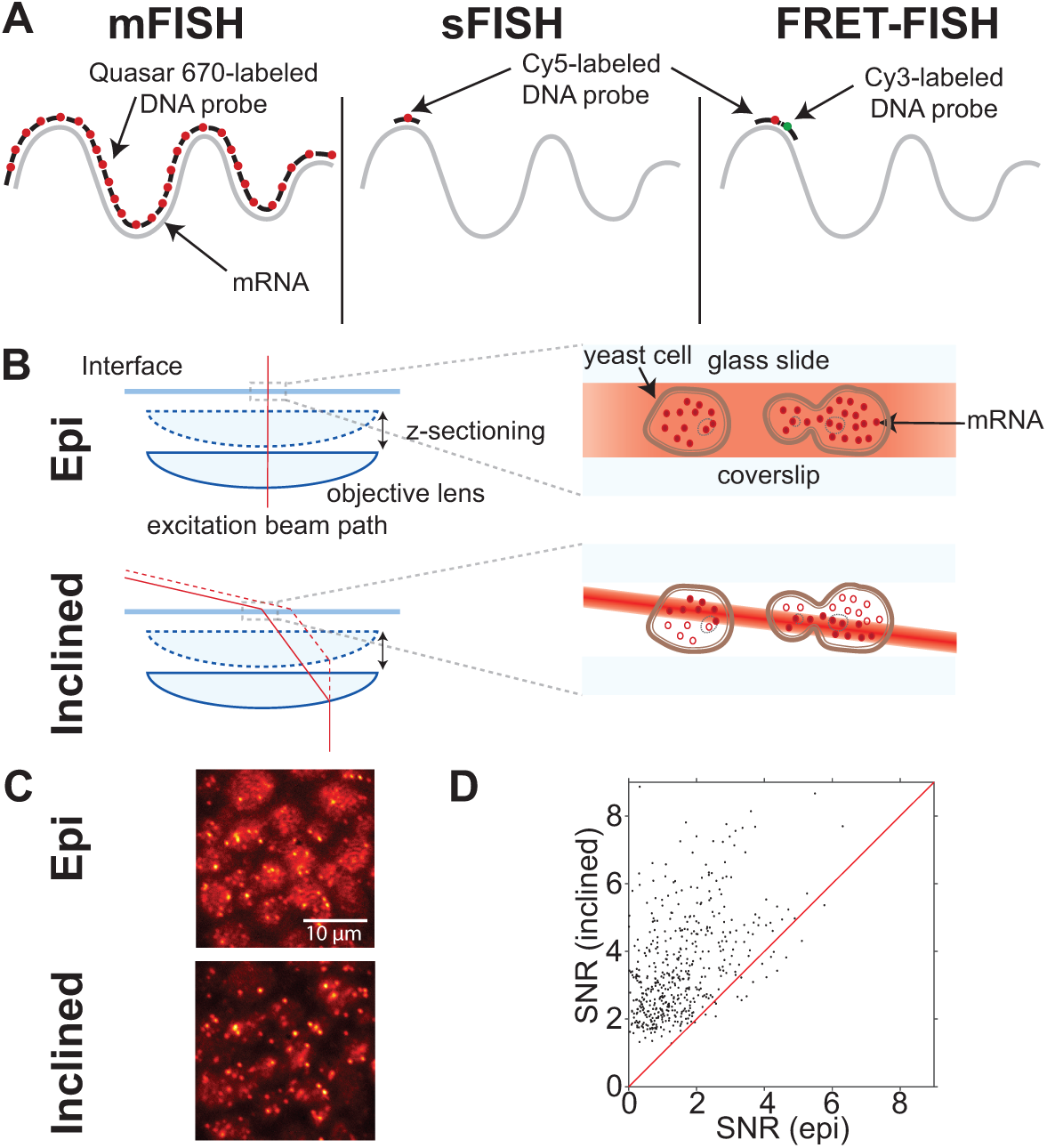
Comparison between single-probe FISH (sFISH) and multi-probe FISH (mFISH). yEVenus mRNA, which is 717-nt long, is probed by sFISH and mFISH. **(A)** Probe configurations are shown from left to right for 30 probe mFISH, sFISH and sFISH with FRET. For mFISH, we use a set of thirty Quasar 670 end-labeled probes. In sFISH, we use a single short Cy5-labeled DNA oligo probe. For FRET experiments the first Cy5-labeled probe is used in conjunction with a Cy3-labeled probe. **(B)** For mFISH, we use a conventional epi-fluorescence microscope setup (top). In epi-illumination, the beam is aligned along the optical axis and illuminates cells across their entire height. The difference in the beam intensity profile between the two geometries is highlighted by varying shades of red. For sFISH, we use highly inclined illumination geometry(bottom), which markedly increases the signal-to-noise ratio (SNR). This light sheet also travels with the objective, which allows imaging different planes along the vertical axis (z-sectioning). **(C)** The images shown are of the same field of view taken with epiillumination (top) and then subsequently with inclined illumination (bottom) using the same laser power. The bottom image taken with inclined illumination exhibits more intense spots and lower background. **(D)** Comparison of spot signal-to-noise ratio (SNR) between epi- and inclined illumination. The SNR values measured with inclined illumination is plotted against those measured with epi-illumination. Most spots are found above the red line *y* = *x*, which indicates inclined illumination produces higher SNR than epi-illumination. The increase in signal to noise ratio is a factor of 2.15 on average.

### B. Correlation between spot count and mRNA level

To test the linearity of our sFISH protocol, we performed FISH on four different strains (Figure 2A) that express yEVenus at four different levels (Supplementary Figure S1). We did not use a promoter inducible by external factors as different induction conditions may differentially affect the hybridization efficiency.

We compared the measured spot counts with yEVenus levels (Figure 2B). The spot count ranged from 2 to 32. We observed a good correlation between spot count and protein level, which argues that fluorescent spots are generated from specific hybridization of the probe to the target mRNA. It also indicates that our spot counting algorithm works well in the range of transcript levels tested.

**FIG. 2.**
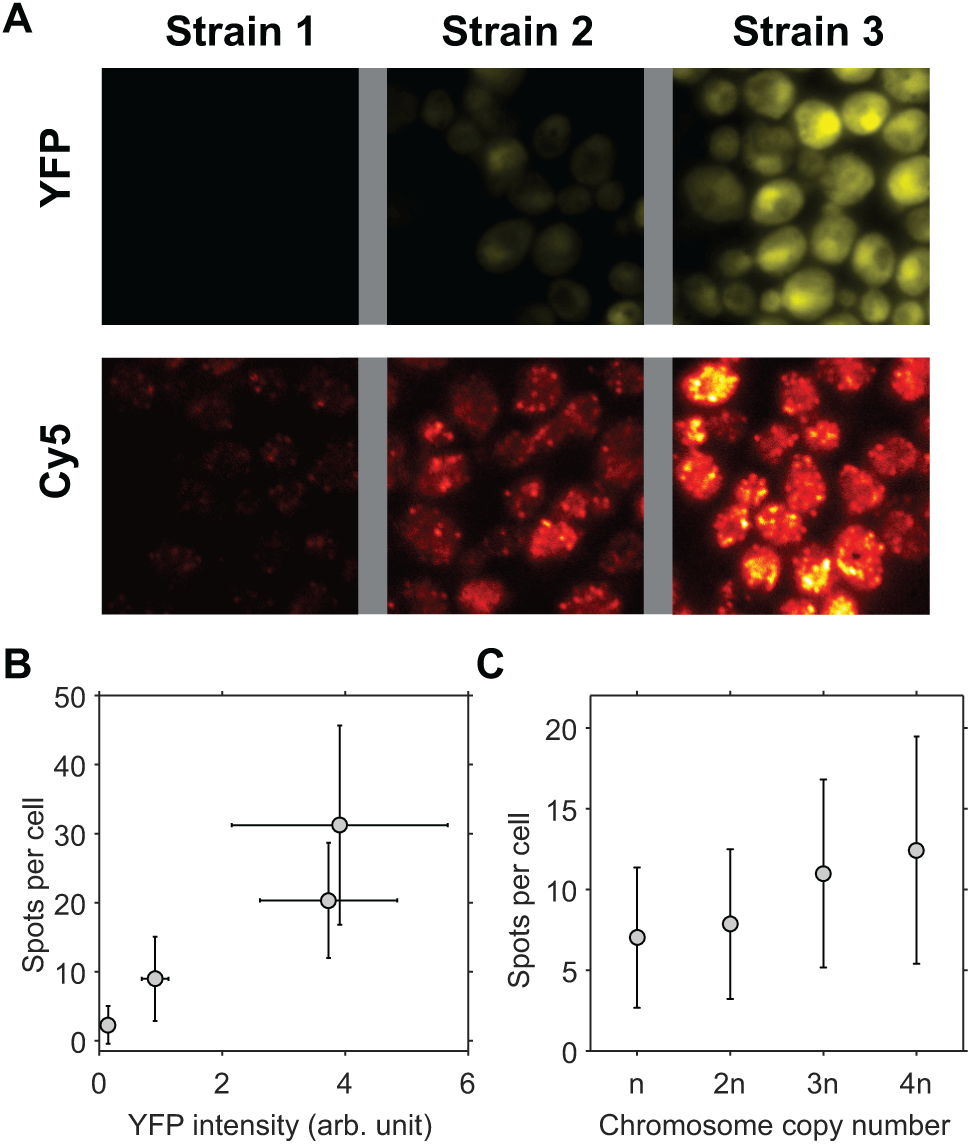
Correlation between sFISH spots and protein expression level. **(A)** Fluorescence images of single yeast cells expressing YFP (top row) and sFISH signals from Cy5-labeled probes targeting YFP mRNA (bottom row). Shown from left to right are fluorescence images of the negative control (no YFP expression), low YFP expression, and high YFP expression. Fluorescence intensities in the YFP channel and Cy5 channel are represented by false yellow and red colors, respectively. YFP images are from formaldehyde fixed cells, and Cy5 sFISH images are from methanol fixed cells. **(B)** Correlation plot. The mean number of FISH spots is plotted vs. the mean yEVenus expression level. The error bars are measures of the standard deviation. **(C)** sFISH spots vs. ploidy. sFISH was performed on yeast strains with four different ploidies (1n, 2n, 3n, 4n). The error bars show the standard deviation of the data. The number of spots detected per cell increases monotonically with the number of copies.

As another control, we performed sFISH on chromosome copy number variants (1n, 2n, 3n, and 4n). We chose to probe a constitutively expressed gene KAP104 (Supplementary Tables S4, S5), which has been used as a reference gene in other studies(35, 36). As expected, the number of spots monotonically increased with the ploidy (Figure 2C), but interestingly, the relationship was not linear, that is, doubling the ploidy did not lead to doubling of the number of spots. This apparent sub-linear relationship could be due to the loss of extra chromosomes or some compensation effect, which will be the subject of future investigation.

In other controls, we were able to mask these sFISH spots with an unlabeled competitor probe in a concentration dependent manner (Supplementary Figure S4C and D), which confirms that the observed sFISH spots are due to the fluorescent probe hybridized to the KAP104 transcript. We also compared sFISH with mFISH (Supplementary Figure S4A). mFISH yielded an average of 9.4 spots per cell, similar to the number previously reported (35, 36). In comparison, sFISH yielded less spots per cell than mFISH (6.5, Supplementary Figure S4B), which suggests that sFISH can detect the KAP104 transcripts with detection efficiency (*p*) of about 64% taking into account the false positive rate (0.5 spots per cell, Supplementary Figure S4D).

### C. Methanol vs. formaldehyde

We initially tried methanol as a fixative following a fast FISH protocol(37), and noticed that fluorescence images looked more clear than when using formaldehyde as the fixative. Hence, we performed a systematic comparison of the two fixatives. We first compared the number of background spots in a negative control strain lacking the yEVenus gene between the two fixatives (Supplementary Figure S8). We found on average 3 spots per cell in formaldehyde-fixed cells compared to 0.3 spots per cell in methanol fixed cells, which indicate that formaldehyde fixation causes more nonspecifically bound or trapped Cy5 probes inside the cell(Figure 3A). Next, we compared the number of spots in a positive control strain with the lowest expression level of yEVenus. In this strain, we detected on average 10 spots per cell with methanol and 3 spots per cell with formaldehyde (Supplementary Figure S7). This result suggests that methanol fixation not only reduces nonspecific binding but also increases the rate of specific binding to the target mRNA.

**FIG. 3.**
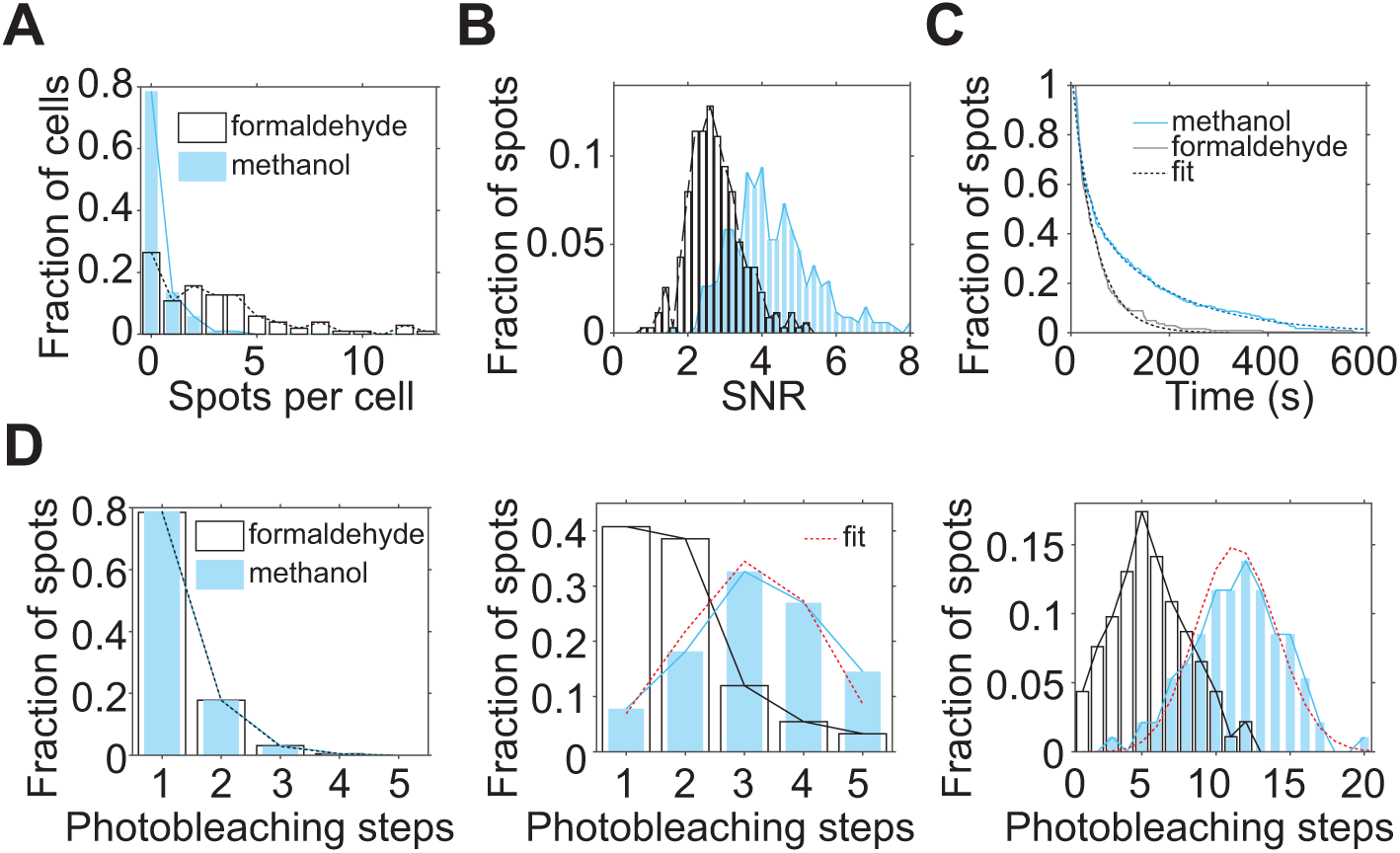
Comparison of spot quality between formaldehyde (white) treated samples and methanol (blue) treated samples **(A)** sFISH spots detected from the negative control strain. On average, there is ~ 0.3 spots per cell in the methanol treated cells (blue) compared to ~ 3.1 spots per cell in the formaldehyde treated sample (white). **(B)** Comparison of signal-to-noise ratio (SNR) of single probes. SNR of a single Cy5 was calculated from fluorescence time traces that captured single-step photobleaching events. Signal is obtained from the single-step drop in fluorescence intensity upon photobleaching, and the noise is calculated as the standard deviation of the signal prior to photobleaching. The histogram shows that the spots from methanol-treated cells (blue) have ~ 2-fold higher SNR than those from formaldehyde-treated cells (white). **(C)** Comparison of Cy5 stability. The population decay curves show that sFISH spots in formaldehyde treated cells photobleach faster than those in methanol-treated cells. **(D)** Comparison of probe number per spot. The number of probes per spot was determined by counting the number of photobleaching steps in the fluorescence time trace. When a single probe was used, most spots photobleached in a single step regardless of the fixative of choice (left). In comparison, when five (middle) or thirty (right) probes targeting the same mRNA were used, more probes were detected from spots in methanol-treated cells than in formaldehyde-treated cells. For the methanol samples treated with multiple probes (middle and right panels), binomial distribution fits are shown in red. For the five-probe experiment (5-probe FISH), Cy5-labeled probes and inclined illumination are used; whereas, for the thirty-probe experiment, Quasar-labeled dyes and epi-illumination are used.

With methanol-fixed cells, we not only detected more spots per cell in the positive control strain, but also detected more fluorophores per spot when multiple probes were used. To count the number of fluorophores per spot, we acquired time series of fluorescence from single spots in a fixed plane until they were completely photobleached. The histograms of the number of photobleaching steps are shown (Figure 3D). When a single probe was used, most spots photobleached in a single step in both methanol and formaldehyde-fixed cells. Spots that photobleached in two steps are likely due to close proximity of different mRNA molecules. As the number of probes was increased (1, 5, and 30), the difference in the number of photobleaching steps between formaldehyde and methanol treated samples became more noticeable. This measurement confirms that methanol fixation allows more efficient hybridization of probes to mRNA while preserving an improved signal-to-noise ratio.

**FIG. 4.**
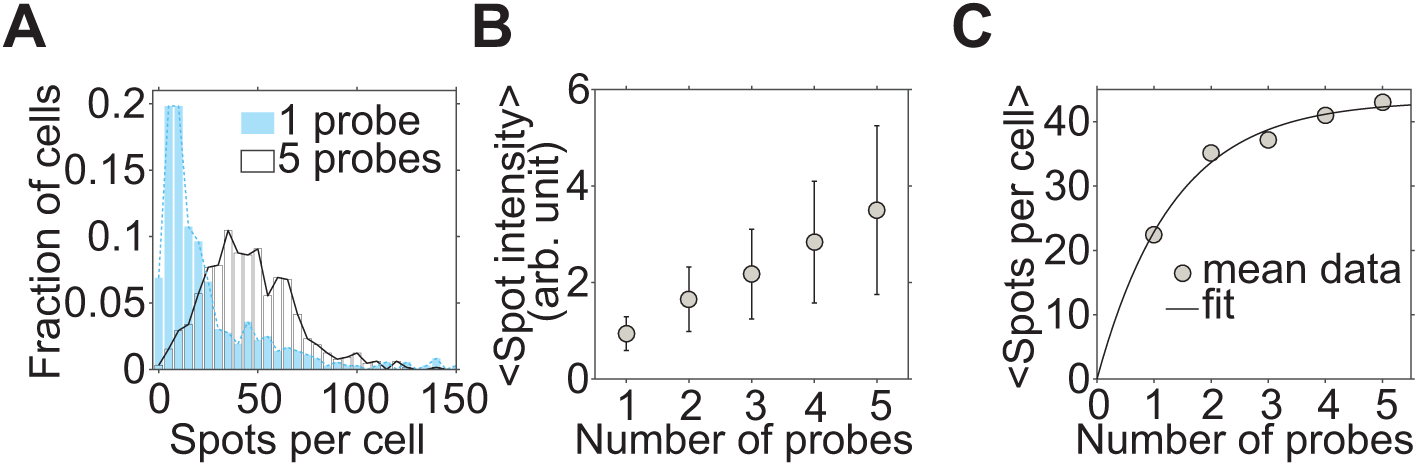
Estimation of the hybridization efficiency of single probes. **(A)** The effect of varying the number of probes. The histograms of the number of spots detected per cell are plotted for 1-probe (blue) and 5-probe (white)FISH. **(D)** Spot intensity vs. probe number. The mean spot intensity increases linearly with the number of probes as expected from the binomial distribution. **(C)** Spot number vs. probe number. The mean number of spots detected per cell (*y*) increases with the number of probes (*x*). The fit model is *y* = *N*(1 – (1 – *p*)*^x^*) where *N* is the true copy number, and *p* is the hybridization rate for a single probe. *p* is extracted to be 53 %.

This data also allows us to estimate the hybridization efficiency of probes. Assuming that probes all hybridize with the same probability *p*, the number of probes per spot can be fitted with a binomial distribution. We fitted the binomial distribution to the two sets of data taken with methanol fixation (red dotted lines, Figure 3D). *p* for yEVenus is extracted to be 61 % for 5-probe FISH, and 38 % for the 30-probe mFISH (Supplementary Table S3). The variation in *p* between the two data could be due to the difference in probe design. In 5-probe FISH, the probes were designed to have similar melting temperatures to the probe used for sFISH (shown in Supplementary Table S2), while in mFISH, the probes are designed to have the same length with no consideration of the melting temperature. Also, poor signal of Quasar 670 used in mFISH can lead to the underestimation of the number of photobleaching steps. Nonetheless, these rough estimates set the detection efficiency in the range of ~40 % to 60 %.

We also characterized some apparent differences in the fluorescence properties of Cy5 due to the difference in fixatives. The fluorescence signal, which is defined as the difference between the fluorescence and background levels, was similar between the two. However, the noise, which is the fluctuation of the Cy5 signal, was significantly higher in formaldehyde fixed cells. As a result, the signal-to-noise ratio was 2-fold higher in methanol-fixed cells(Figure 3B). In addition to having an advantage in signal to noise ratio, methanol treated cells exhibited a longer Cy5 lifetime (Figure 3C).

### D. Detection Efficiency

To ensure that our FISH protocol operates at maximum hybridization efficiency, we increased zymolyase incubation time or the probe concentration until the spot count did not increase further (Supplementary Figures S9, S10, and S11). Even under this condition, however, our single-probe protocol is expected to underestimate the actual number of mRNA transcripts due to hindered accessibility of the target region of some transcripts. We can also estimate the effective detection efficiency (*p*) for the yEVenus transcript by increasing the number of probes. The number of spots detected per cell initially increased with the number of probes, but soon plateaued at four to five probes (Supplementary Figure S6). Assuming that each probe binds the target mRNA with probability *p*, the probability of failing to detect an mRNA molecule with n probes is (1 − *p*)*^n^*. Therefore, the number of detected spots (*y*) should increase with the number of probes (*x*) as *y* ∝ 1 − (1 − *p*)*^x^*. We fitted this model to the plot of the spot count per cell vs. probe number (Figure 4) and extracted p for the yEVenus transcript to be 53 %, which is consistent with the range determined in the previous analysis (Figure 3).

**FIG. 5.**
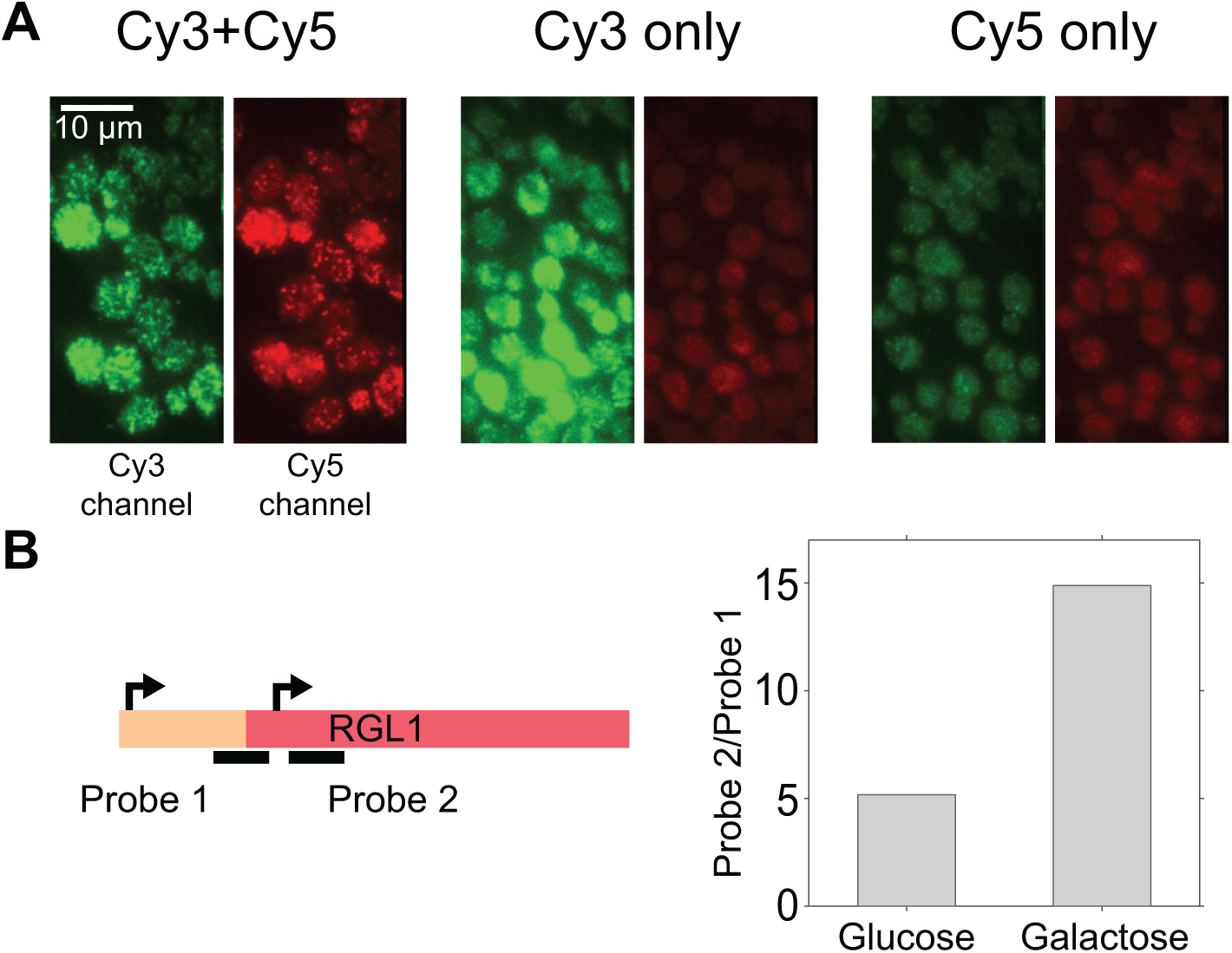
Applications of sFISH. **(A)** Demonstration of FRET-FISH in yeast. Fluorescence image acquired under 532-nm excitation was split into two half images based on the emission wavelength. In each image, the green half on the left is from the Cy3 emission channel, and the red half on the right is from the Cy5 emission channel. The images shown represent cells treated with both Cy3- and Cy5-probes (left), Cy3-probe only (middle), and Cy5-probe only (right). Bright, punctate spots were observed in the Cy5 channel only when cells were treated with both probes (left). **(B)** sFISH for mRNA isoform detection. The schematic on the left depicts alternative transcription initiation sites (arrows) at the RGL1 locus, which lead to mRNA isoforms with different lengths. Transcription from the first site produces a full-length mRNA, while from the second site produces a truncated isoform. Using sFISH with two separate probes, the relative fractions of these isoforms can be measured. Probe 1 targets the longer isoform only, whereas probe 2 targets both. The bar plot on the right shows the ratio of sFISH signals with probe 2 to probe 1 measured with glucose (left) or galactose (right) growth media. Here, the mean total fluorescence intensity per cell was used as a proxy for sFISH signal because transcription level was too high to count individual spots.

### E. mRNA detection via FRET

In addition to various methods used above to infer the detection efficiency *p*, we also tried to determine *p* using Förster Resonance Energy Transfer (FRET). In this approach, two DNA probes complementary to immediately adjacent regions of the mRNA are labeled with donor (Cy3) and acceptor (Cy5), respectively, so that the FRET signal would arise only when both probes bind to the same mRNA (Figure 1A). By comparing the number of fluorescent spots due to FRET to the number of fluorescent spots due to direct excitation, the detection efficiency can be directly determined.

sFISH was performed with donor and acceptor probes at 1:1 ratio at the same concentration used for other experiments. As controls, sFISH was also performed while leaving out one of the probes. The typical sFISH images from three different combination of probes are presented in Figure 5. Upon 532 nm excitation, signal in the Cy5 channel was visible only when Cy3 probe is present (left, Figure 5), which indicates that many mRNA molecules are hybridized with both the Cy3-probe and Cy5-probe. We confirmed that this intense Cy5 signal could not have resulted from bleedthrough of Cy3 signal into the Cy5 channel (middle, Figure 5) or direct excitation of Cy5 by the 532 nm laser (right, Figure 5). Upon 532-nm excitation, spots that appear in the Cy5 channel are due to FRET from the Cy3-probe to the Cy5-probe bound to the same mRNA. On the other hand, spots in the Cy3 channel arise from mRNA molecules bound with the Cy3-probe only. We can thus estimate the detection efficiency by dividing the number of Cy5 spots by the total number of both Cy3 and Cy5 spots. Using this method, the detection efficiency is determined to be 48 %.

### F. mRNA isoform detection via sFISH

Since sFISH requires only a 20-30 nt RNA target, it can be used to differentiate mRNA isoforms that are only slightly different in length or sequence, thus offering more versatility than mFISH. As a proof of principle, we chose gene RGL1 (YPL066W), which exhibits differential usage of alternative transcription sites between glucose and galactose growth media(38) (Supplementary Figure S13). As shown in the simplified schematic in Figure 5B, initiation normally starts upstream of the open reading frame (ORF) of RGL1 and produces a full-length transcript, but it can also start within the ORF and produce a truncated isoform. To measure the isoform profile, we designed a pair of probes (Supplementary Figure S13, Supplementary Table S6) that flank the truncation site (solid lines, Figure 5B) and performed sFISH with each probe on yeast cells grown in glucose and galactose media. Since the transcription levels were too high for reliable spot counting, we instead used the total fluorescence intensity integrated over the volume of the cell as a proxy for the transcription level. In qualitative agreement with the genome-wide transcript isoform study(38), we found that the truncated isoform is significantly enriched over the full-length isoform in galactose-containing media (Figure 5B).

## IV. DISCUSSION

In this work, we show that mRNAs in single yeast cells can be quantified using only a single ~20-nt probe labeled with a single Cy5 dye. In this new yeast FISH protocol, we substantially improved signal-to-noise ratio and detection efficiency compared to current protocols by using a combination of methanol fixation and inclined illumination. We measured the detection efficiency in situ based on (i) the number of spots obtained with sFISH and mFISH, (ii) the number of photobleaching steps per fluorescent spot, (iii) the number of fluorescent spots per cell, and (iv) the number of fluorescent spots due to FRET. These methods yielded detection efficiencies above 50 % with some variability possibly due to the difference in probe sequence and length.

To detect single Cy5-labeled probes in situ, we used a lab-built fluorescence microscope with single molecule sensitivity. In this setup, 640-nm laser is focused off-center at the back focal plane of a high NA objective to obtain highly inclined illumination. Inclined illumination excites a smaller volume compared to epi-illumination, leading to a higher signal and lower background. We expect more advanced light-sheet microscopy setups (39–42) to perform equally well or better but they would require a specialized sample chamber for side-illumination. In principle, imaging flow cytometers with an extended depth of field can be used to increase throughput for counting fluorescent spots (43). Whether their sensitivity is sufficient to measure sFISH spots needs to be tested in the future.

We used backbone-integrated Cy5, which has a higher photostability(14) than base-linked Cy5 used in previous studies. Photobleaching lifetime of Cy5 is extended into minutes using PCA-based oxygen scavenging(29). Since the PCA-based system does not have heme groups or flavins, it also exhibits lower autofluorescence than glucose-based oxygen scavenging system.

Methanol, which is the fixative used in sFISH, perforates the cell membrane(44) by removing the phospolipids(45). Hence, the methanol-based method requires less stringent zymolyase treatment for probe delivery compared to formaldehyde-based method. The optimal digestion condition is important for accurate quantification of mRNA level; underdigestion leads to false negatives whereas overdigestion leads to leakage of cytoplasmic material including mRNA. However, we find it difficult to generate a uniformly digested population of cells using zymolyase only. In comparison, the combination of methanol treatment with mild zymolyase treatment produces uniformly permeable population of cells in a reproducible manner.

In contrast to the cross-linking agent formaldehyde, methanol also induces denaturation of cellular proteins(46) including mRNA binding proteins (47), which may explain the increase in hybridization efficiency(48) and reduction of cellular autofluorescence(49, 50). In comparison, formaldehyde is known to modify the amine group available in nucleic acids(51), most notably in the guanine base(52), which will inevitably compromise hybridization efficiency. Moreover, we observed that formaldehyde fixation gave rise to higher background.

Our sFISH method can detect up to 64 % of mRNA transcripts (See Supplementary Figure S4D), which is similar to the detection efficiencies reported in other FISH studies (61 % in budding yeast(20, 53), 65%(11) in human melanoma cells, and 70%(54) in CHO cells). The detection efficiency extracted from our in situ analysis is also consistent with the fact that our sFISH yields about half as many spots as mFISH does for both the PHO5 mutant(24, 55, 56) and KAP104(36).

Spot counting becomes difficult when the transcript level is high because spots begin to overlap with one another. In such cases, the total fluorescence intensity from a single cell can be used to extrapolate the transcript level(55, 57, 58). Compared to mFISH, our sFISH produces spots with more uniform intensities because mRNA is associated with a single fluorophore, and therefore, the total fluorescence intensity would be a robust proxy for the transcript level.

Our sFISH method not only offers a time- and cost-efficient mRNA quantification tool, but also opens up new opportunities for transcript research at the single cell level. Owing to its much smaller target size compared to mFISH, sFISH can be used to detect RNA shorter than 100 nt, profile mRNA isoforms (38, 59, 60), detect subtle changes in mRNA sequence over a 30-nt window as a result of alternative splicing (61>–63) and to measure transcriptional(64) and degradational intermediates (65) with improved resolution. FRET-FISH, which we demonstrated here, can also be used to detect circular RNA (66) in situ without an amplification step.

## V. CONCLUSION

We demonstrate sFISH to be viable for budding yeast cells. Through the combination of highly inclined illumination and methanol fixation, improved accuracy and consistency in quantification of mRNA level can be achieved. This method also yields comparable results to mFISH protocol at a reduced cost. We envision this method to be particularly useful for quantification of a short stretch of RNA at the single cell level.

## VI. ACKNOWLEDGEMENTS

This work is supported by Georgia Institute of Technology startup funds, GAANN Molecular Biophysics and Biotechnology Fellowship, and the National Institutes of Health grant (R01-GM112882). J.S.C. thanks the Litovitz Family Fund for support.

---

**Supplementary Figure S1.**
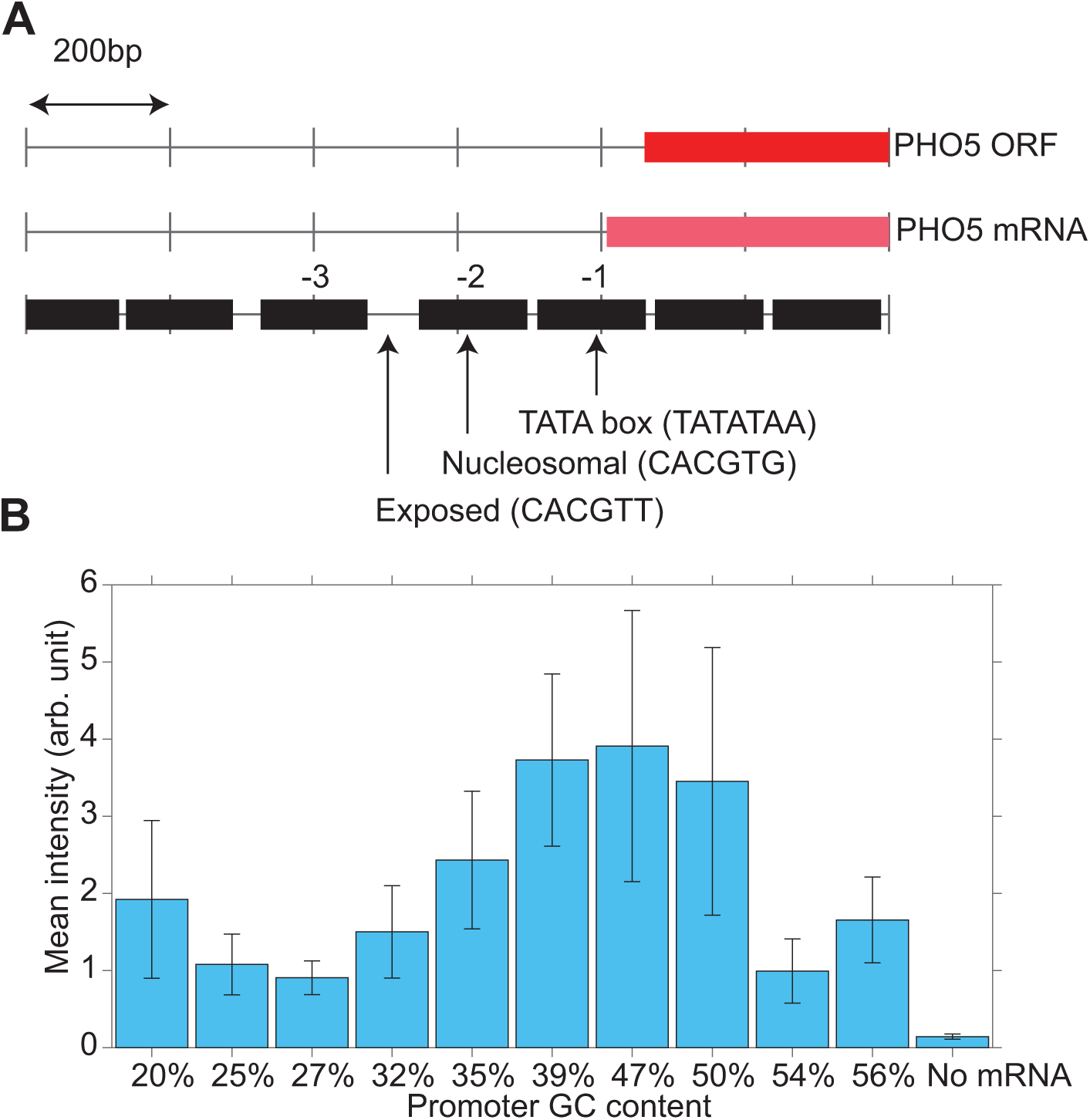
PHO5 promoter variants used in this study. (A) PHO5 promoter map. The open reading frame (ORF) and the transcribed region of the PHO5 gene are shown. The reference nucleosome map is retrieved from (67). The first three nucleosomes are numbered from -1 through -3. The wild-type PHO5 promoter contains two Pho4 binding sites, one in the exposed region between nucleosome -3 and -2, and one within nucleosome -2. The nucleosomal site (CACGTG) has a stronger affinity to Pho4 than the exposed site (CACGTT). The promoter variants used in this study have a common lone high affinity site (CACGTG) in the exposed region with various GC% sequences in the nucleosome -2 region. (B) YFP Expression Level. YFP intensity is plotted against the percentage of GC in the promoter sequence of the strain. This is a non-monotonic relationship that shows the lowest expression in the 27%GC strain and the highest in the 47%GC strain. Error bars show the standard deviation of the population which is much larger for strains that are more highly expressing.

**Supplementary Figure S2.**
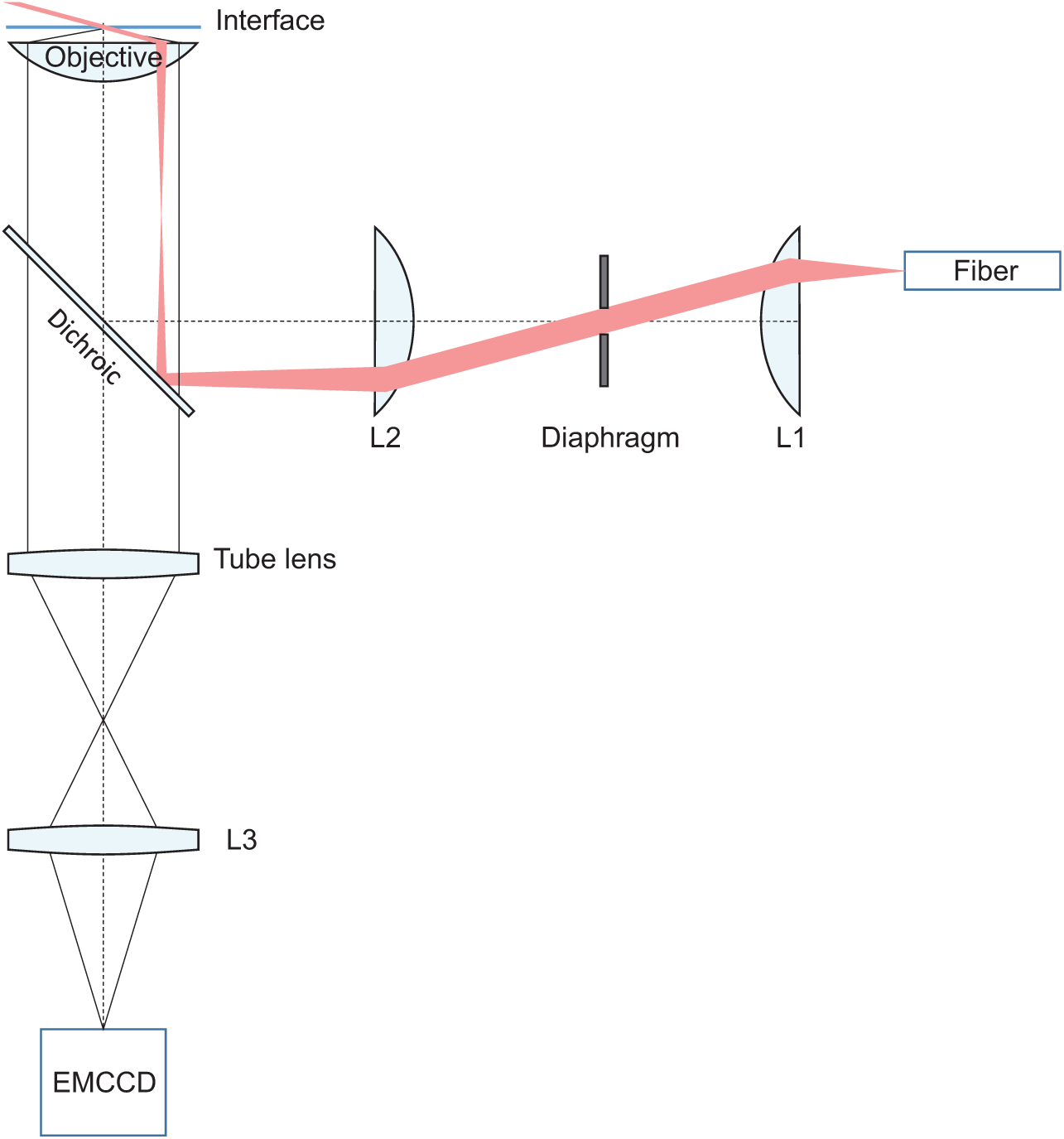
Optical setup for fluorescence microscopy. A laser line is selected by the AOTF (not shown) and coupled to the fiber optic. The fiber output can be laterally translated with respect to lens L1 to vary the incidence angle. L1 collimates the output beam from the fiber, and lens L2 focuses the beam on the back focal plane of the objective. The illumination area is adjusted using the diaphragm located at the image conjugate plane between L1 and L2. The image formed by the tube lens (f=18 cm, Olympus) outside the microscope is relayed onto the EMCCD by lens L3 with some magnification.

**Supplementary Figure S3.**
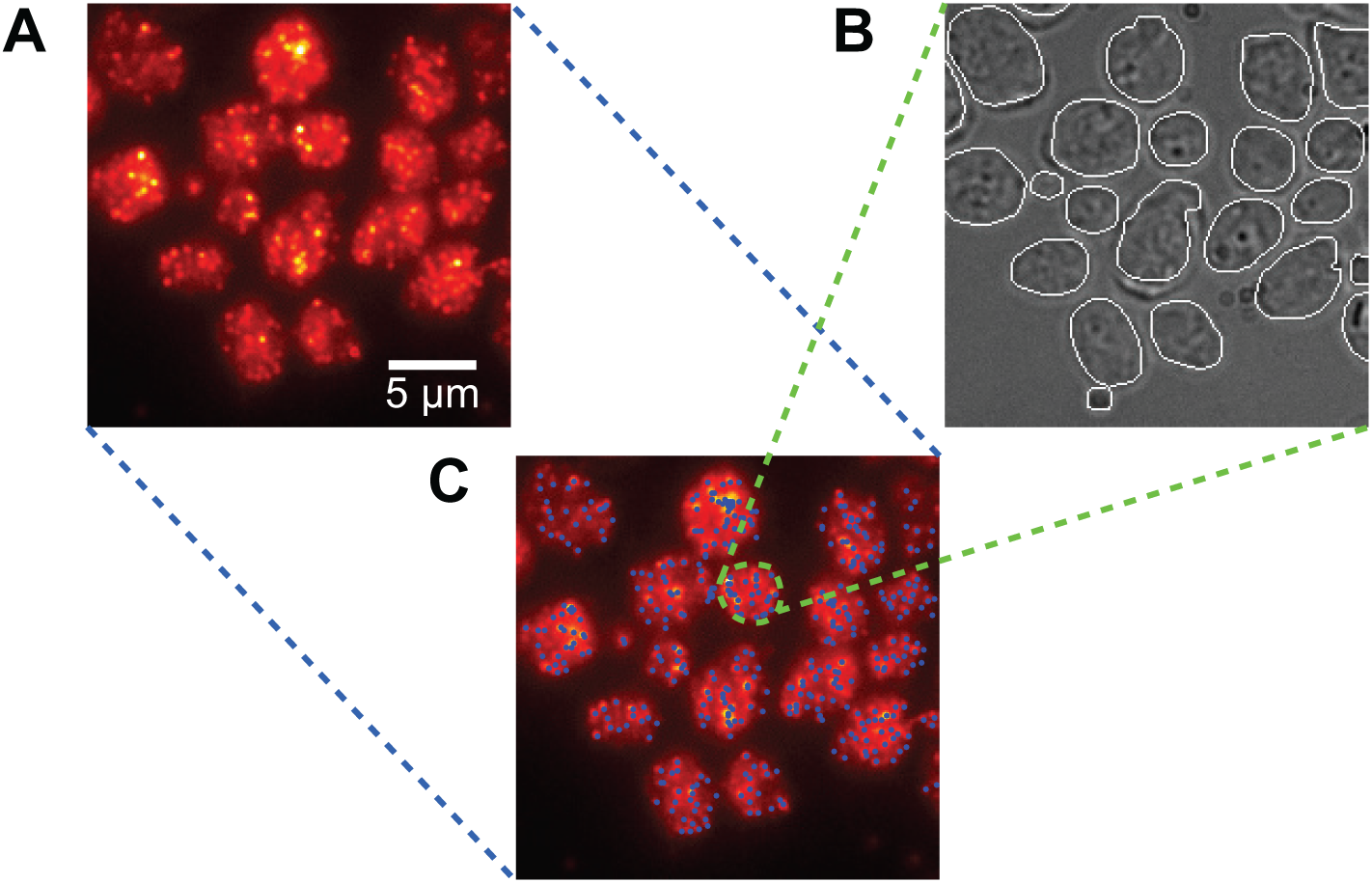
Image processing. (A) Raw sFISH data from the Cy5 channel for an mRNA expressing strain. (B) Detection of cell boundaries by applying Sobel filter on the DIC image stack. The local background is approximated by averaging the pixels near the boundary. (C) Spot detection. All local maximum intensity pixels are considered candidate spots. The distribution of the background-subtracted spot intensities exhibits a peak near zero and another peak centered at a higher intensity. Only the spots that belong to the higher intensity peak are qualified as true spots.

**Supplementary Figure S4.**
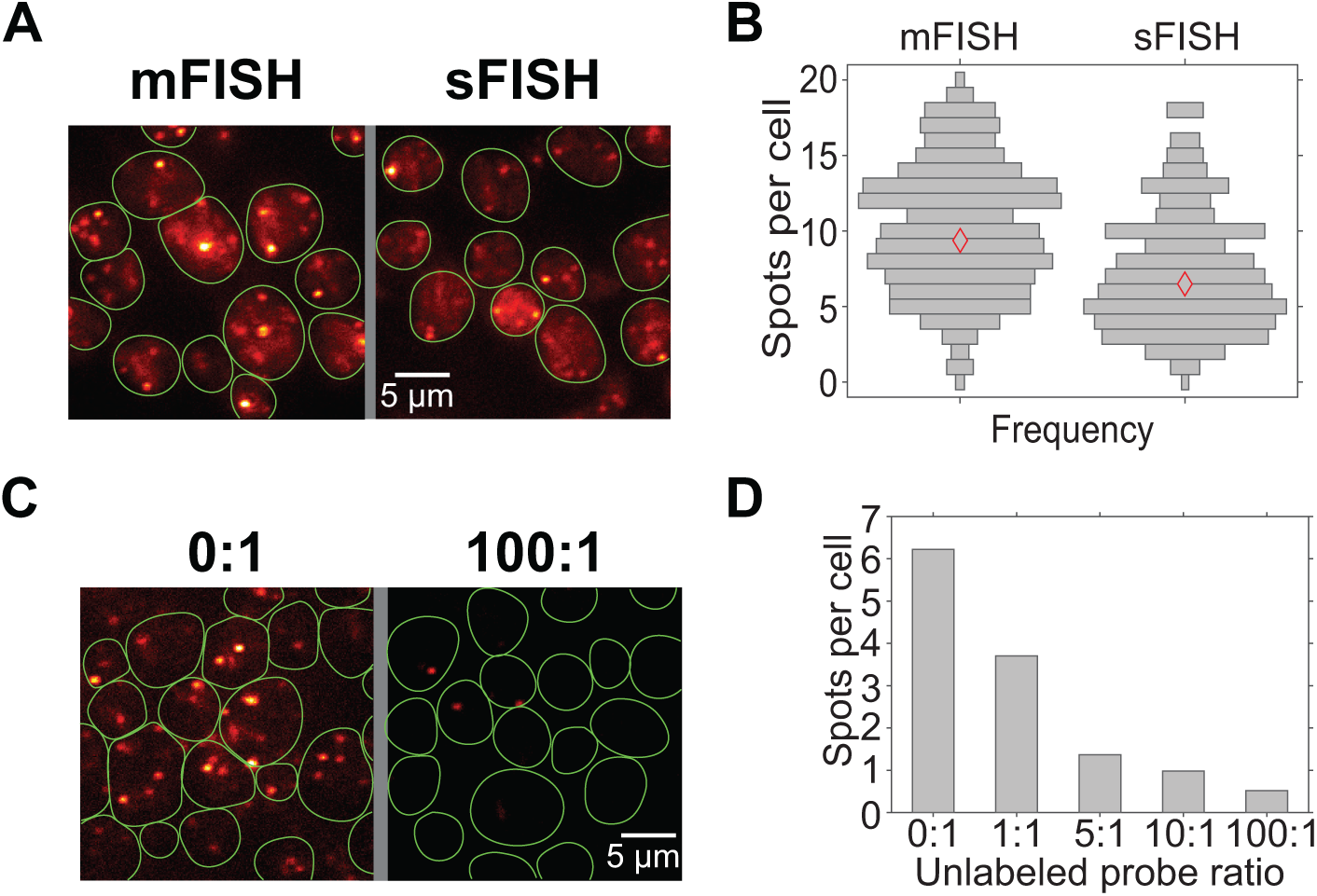
Control experiments with KAP104 FISH. (A) Comparison between mFISH and sFISH. mFISH and sFISH targeting the KAP104 transcript were performed on the cells grown in the same tube, but fixed with either formaldehyde (mFISH) or methanol (sFISH). Cell boundaries are shown in green. In both cases punctate spots can be seen. mFISH spots are only 2 to 3 times brighter than sFISH spots despite using as many as 48 probes. The relatively low fluorescence signal of mFISH spots compared to sFISH spots is due to the dye (Quasar 670 vs. Cy5) and illumination geometry (epi vs. inclined). (B) Spot count distributions from mFISH and sFISH. In these horizontal bar graphs, the position on the vertical axis represents the number of spots per cell, and bar width represents the frequency. Red diamonds are the mean values (9.4 for mFISH and 6.5 for sFISH). The wide distributions represent cell-to-cell variability, not experimental noise. (C) sFISH images with (right) and without (left) an unlabeled competitor probe. The ratio above each image is the ratio of unlabeled to labeled probes. (D) Mean number of spots per cell vs. unlabeled probe. The spot count decreases monotonically with the concentration of unlabeled probes. The concentration of labeled probes is fixed in the protocol. At 100:1 ratio, only ~0.5 spots per cell are seen, which is similar to the rate of false positives.

**Supplementary Figure S5.**
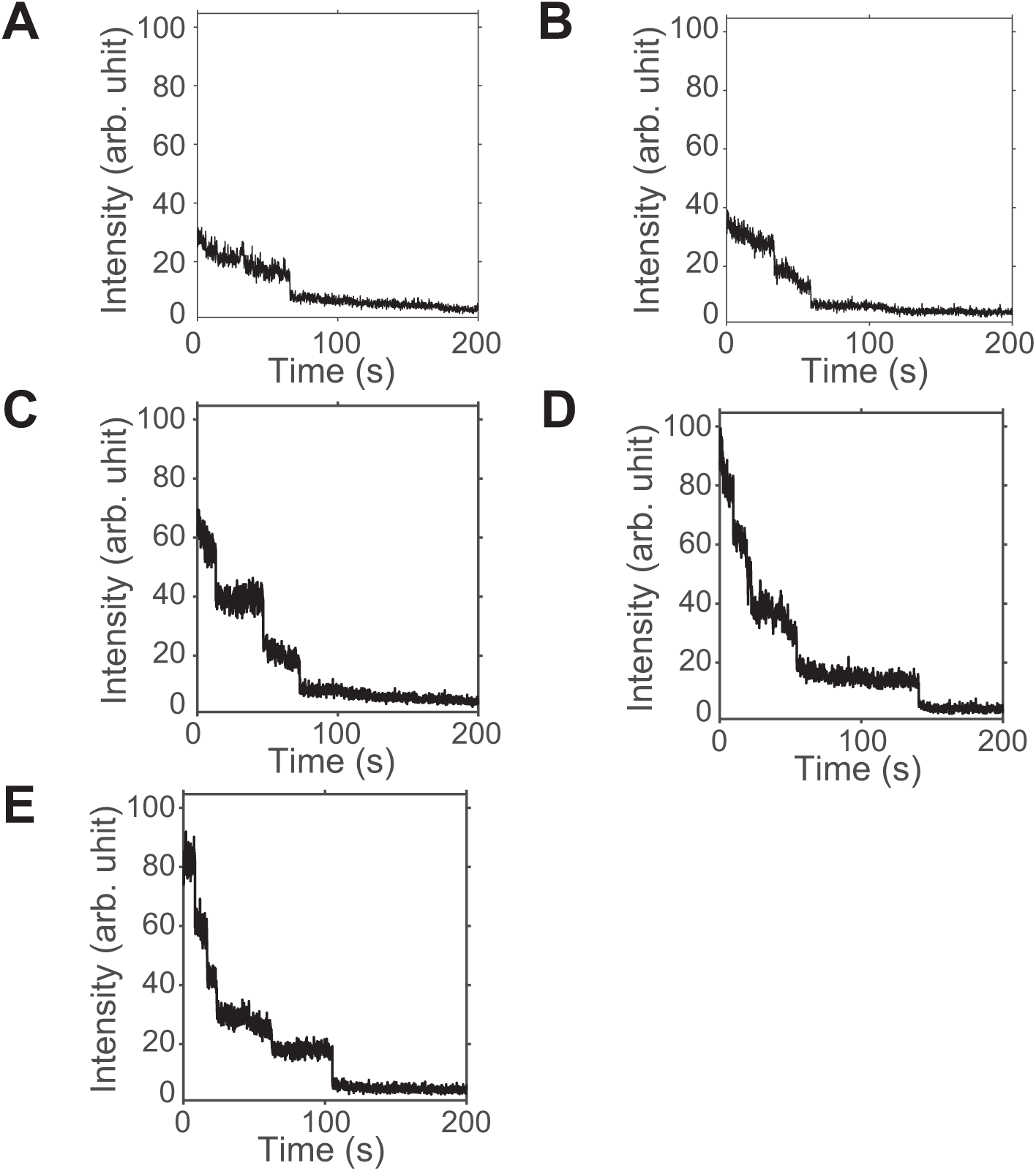
Photobleaching of FISH spots. Fluorescence intensities from single spots were monitored under continuous excitation. Most sFISH spots show photobleaching in a single step, (A), which is evidence for the presence of a single fluorophore. Subsequent panels from (B) to (E) show traces taken from 2, 3, 4, and 5 probe treatments, respectively. Overall, the number of photobleaching steps increases with the number of probes used. For these acquisitions, the exposure time was set to 100ms.

**Supplementary Figure S6.**
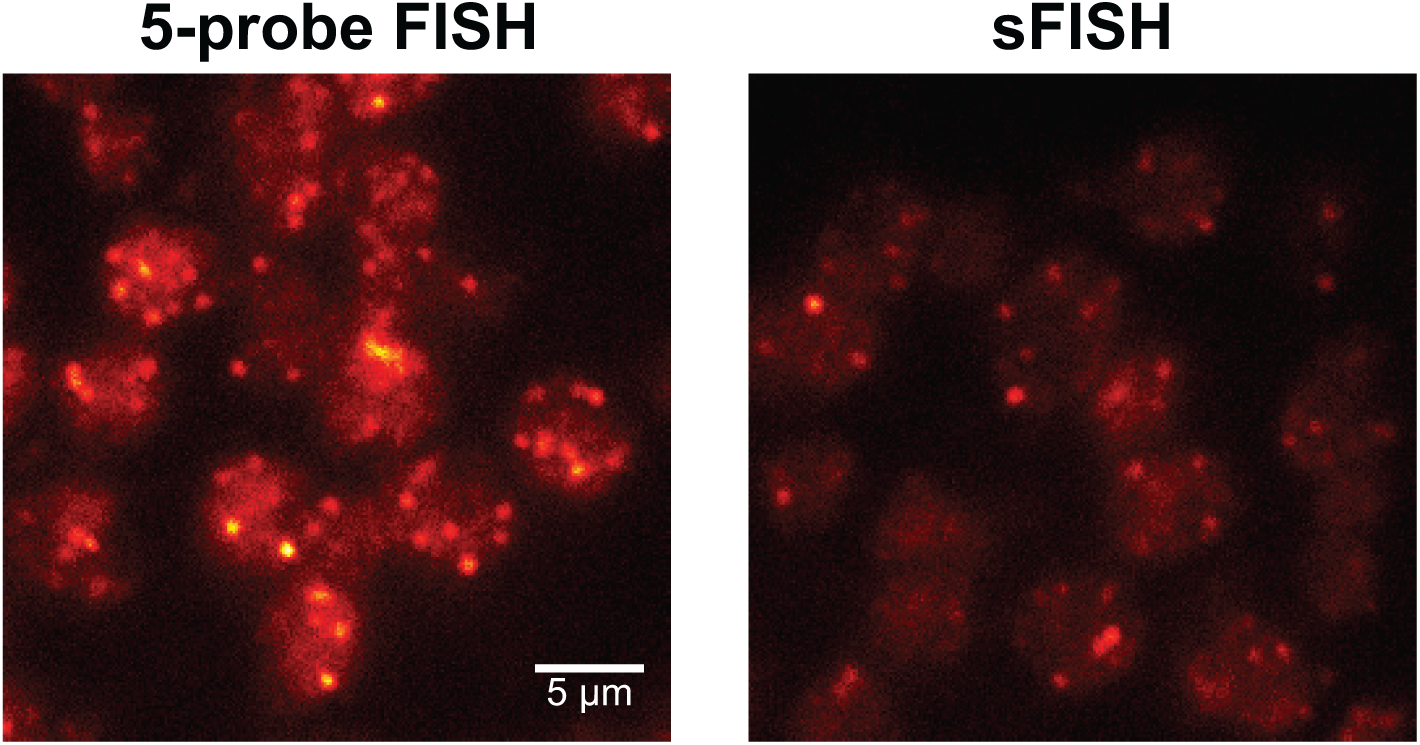
Comparison of FISH with five probes and a single probe. Raw FISH images for the low expression yEVenus strain are shown for five Cy5-labeled probes (left) and a single Cy5-labeled probe (right). The average intensity of spots is about three-fold higher when five probes are used.

**Supplementary Figure S7.**
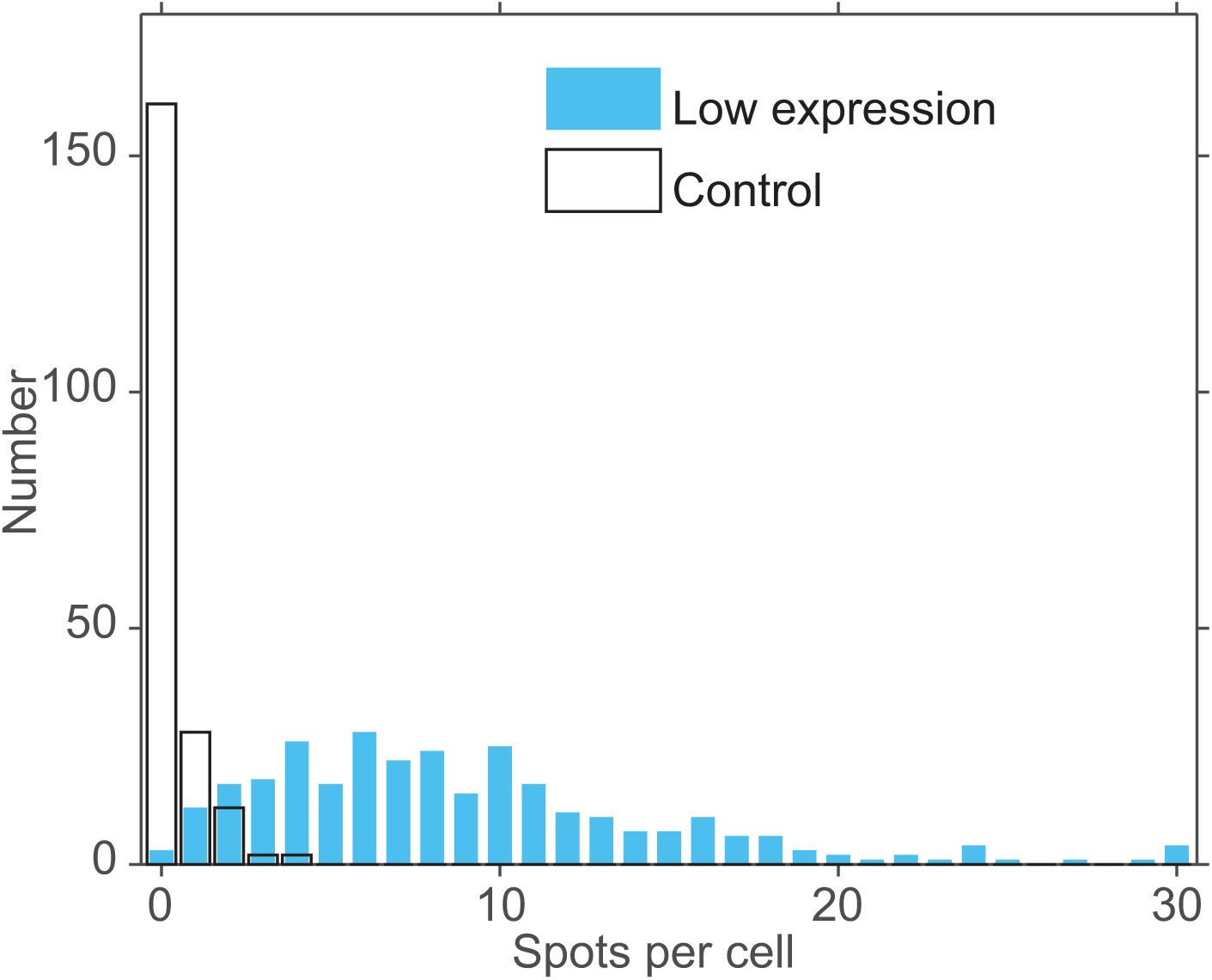
False positives in sFISH. Spots counted in each cell are plotted as a histogram. The negative control strain yields a false positive rate of less than 1 per cell (transparent bars). For comparison, the distribution from the low expression strain (positive control) is shown in blue bars.

**Supplementary Figure S8.**
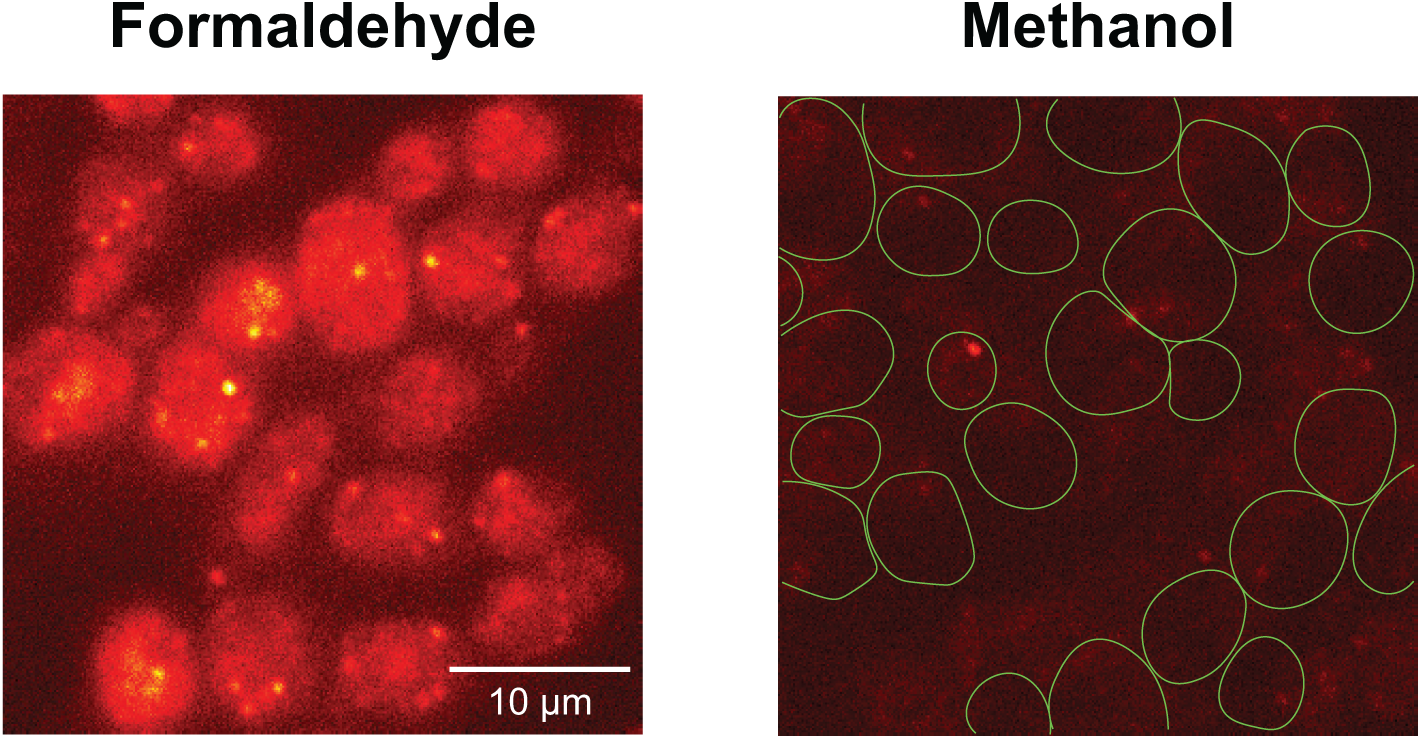
Formaldehyde vs. methanol. Negative control cells treated with single probes are shown with formaldehyde fixation (left) and methanol fixation (right). Formaldehyde-fixed cells exhibit higher cellular background as well as more punctate spots (false positives) than methanol-fixed cells. Cell boundaries are shown only for methanol-treated cells.

**Supplementary Figure S9.**
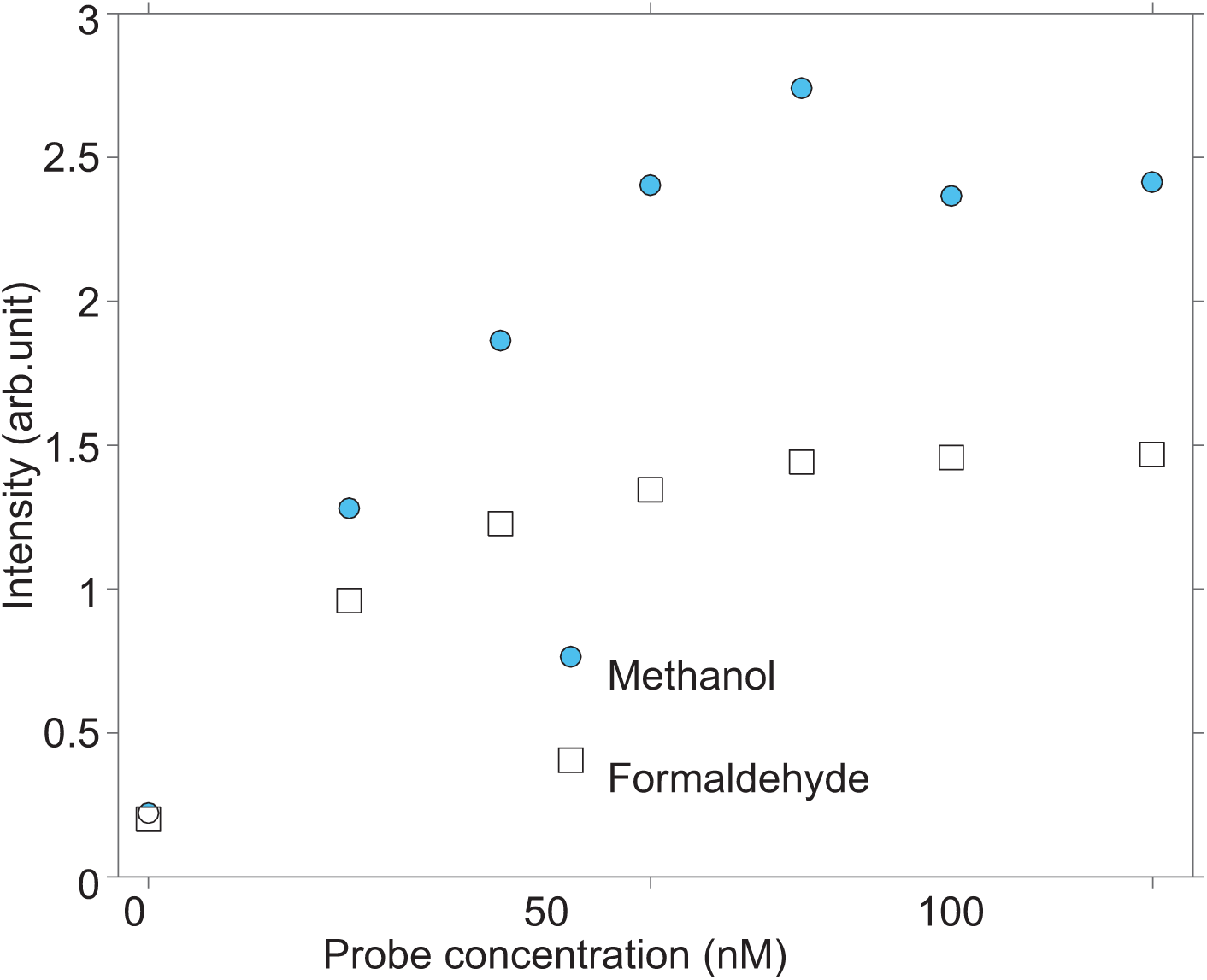
sFISH signal vs. probe concentration. sFISH was performed on both methanol- and formaldehyde-fixed cells over a range of probe concentrations. The plot shows that the average fluorescence signal per cell increases with probe concentration and plateaus around 60 nM. Each data point is an average from 200-300 cells. Based on this relationship, the probe concentration of 65 nM is selected for the standard sFISH protocol.

**Supplementary Figure S10.**
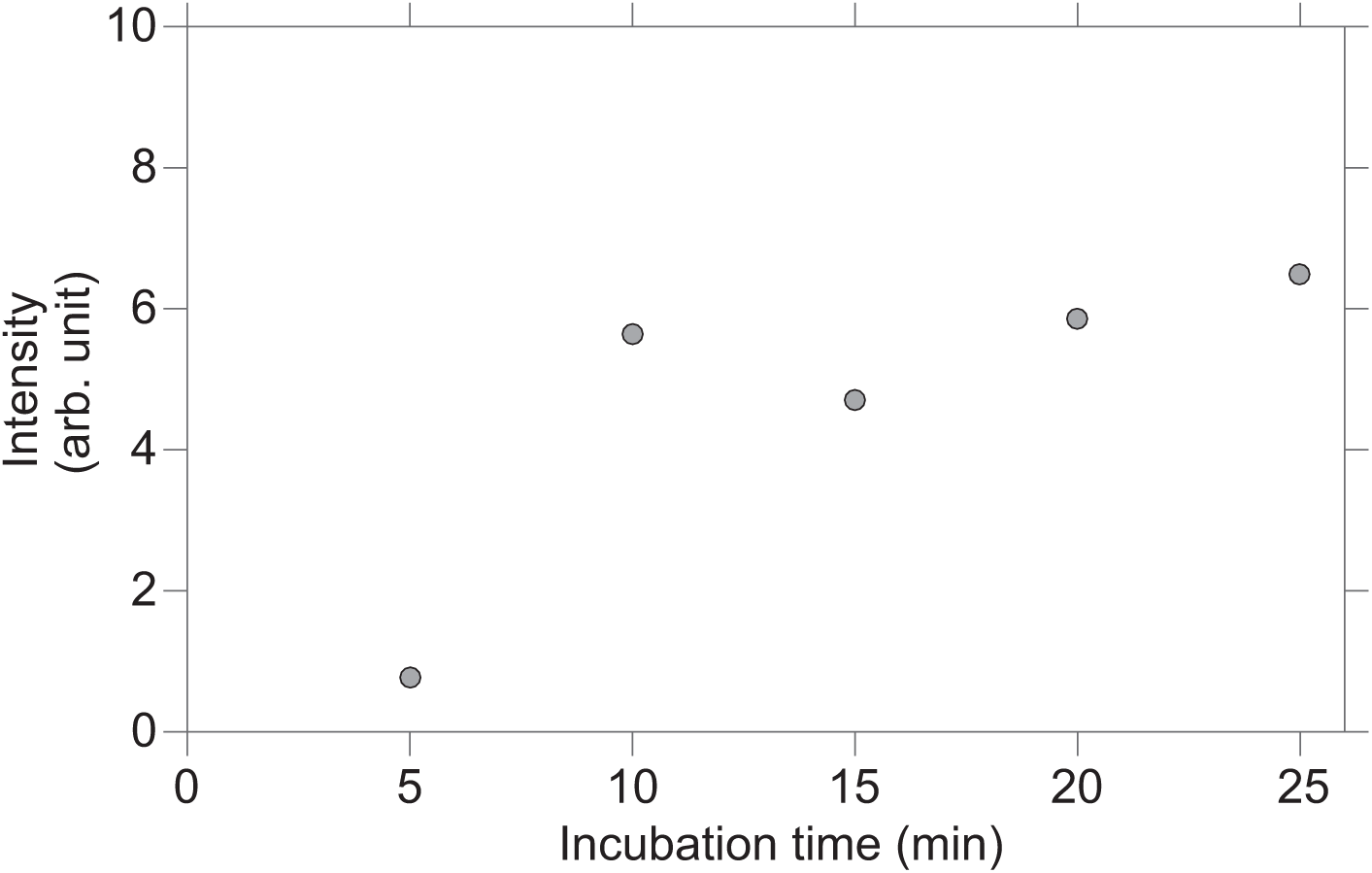
sFISH signal vs. zymolyase incubation time. sFISH was performed on cells spheroplasted in zymolyase for different amounts of time. The subsequent probe treatments were identical. As shown in the plot, the average fluorescence intensity per cell plateaus at 10 minutes of incubation. Each data point is an average from 200-300 cells.

**Supplementary Figure S11.**
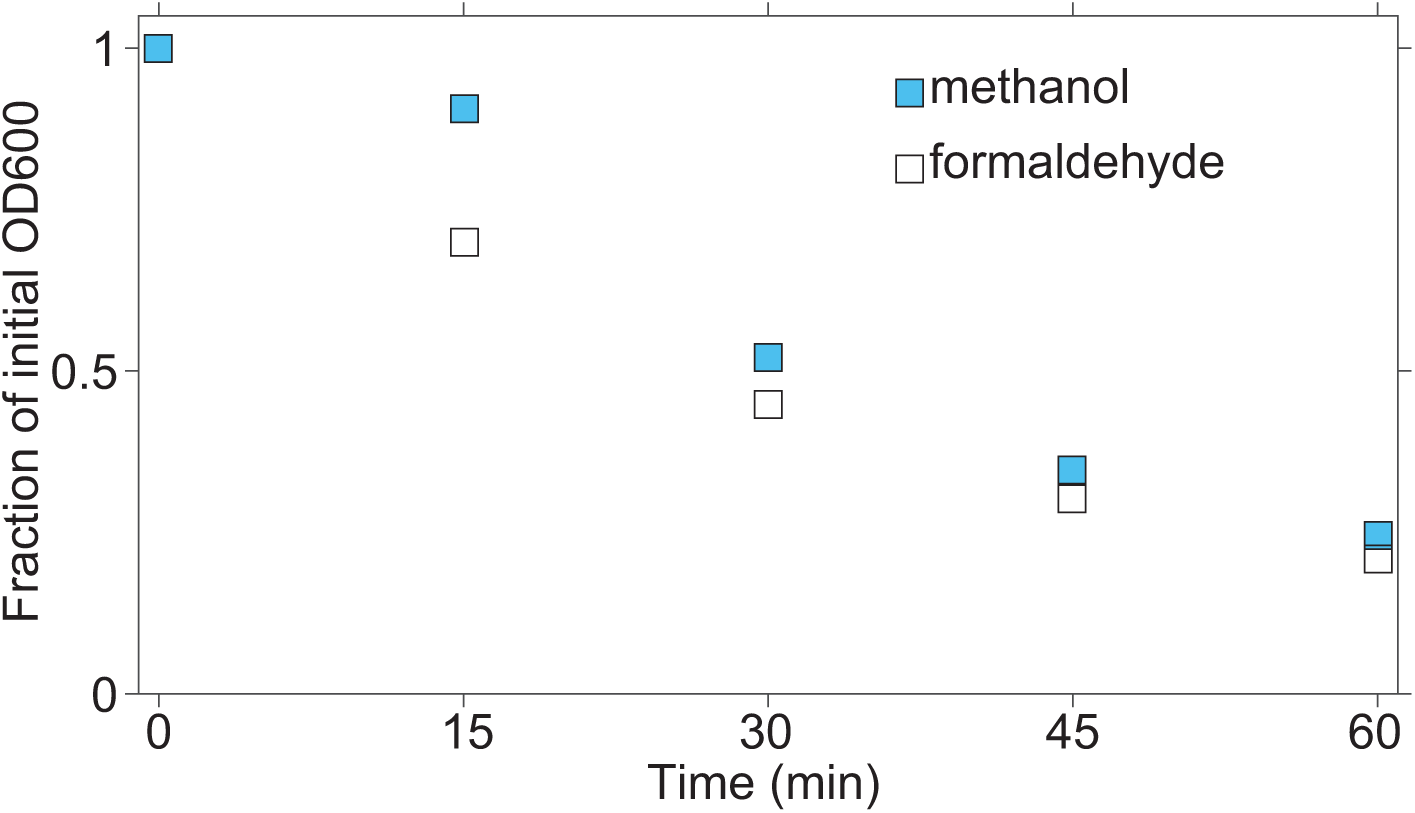
Spheroplasting by zymolyase confirmed by absorbance measurement (OD600).

**Supplementary Figure S12.**
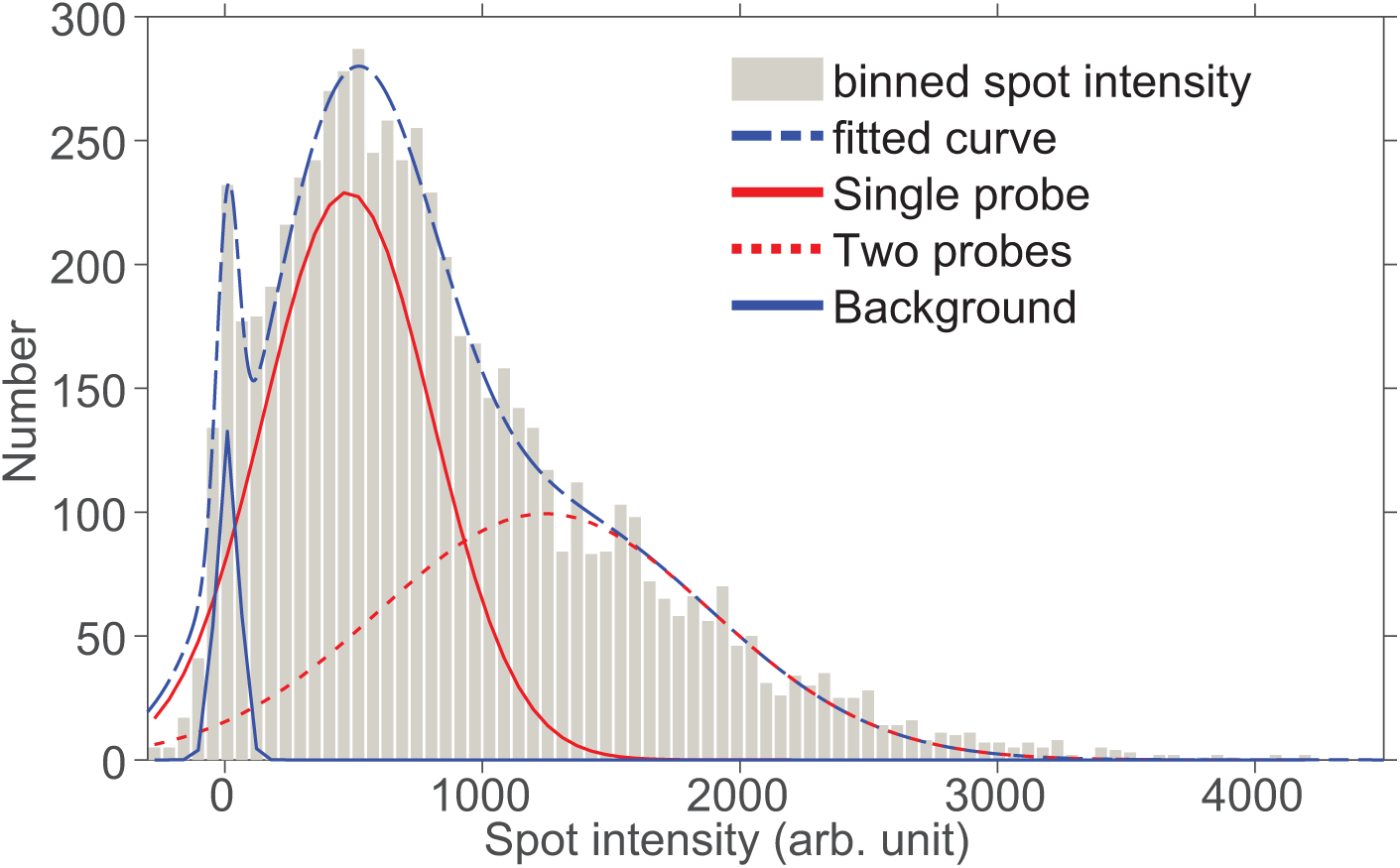
Spot ambiguity. Spots inside a cell are not uniformly bright. Using a single fluorophore requires every spot to be considered equally. By integrating the area under the highest intensity Gaussian we can say how often an ambiguous spot occurs. In our lowest expressing cell we find on average one ambiguous spot. We can say that the fit represents background peaks, single fluorophores and ambiguous spots. We find that the rate of ambiguous spots in our highest expressing strain is about 4.4 per cell (13% of spots.)

**Supplementary Figure S13.**
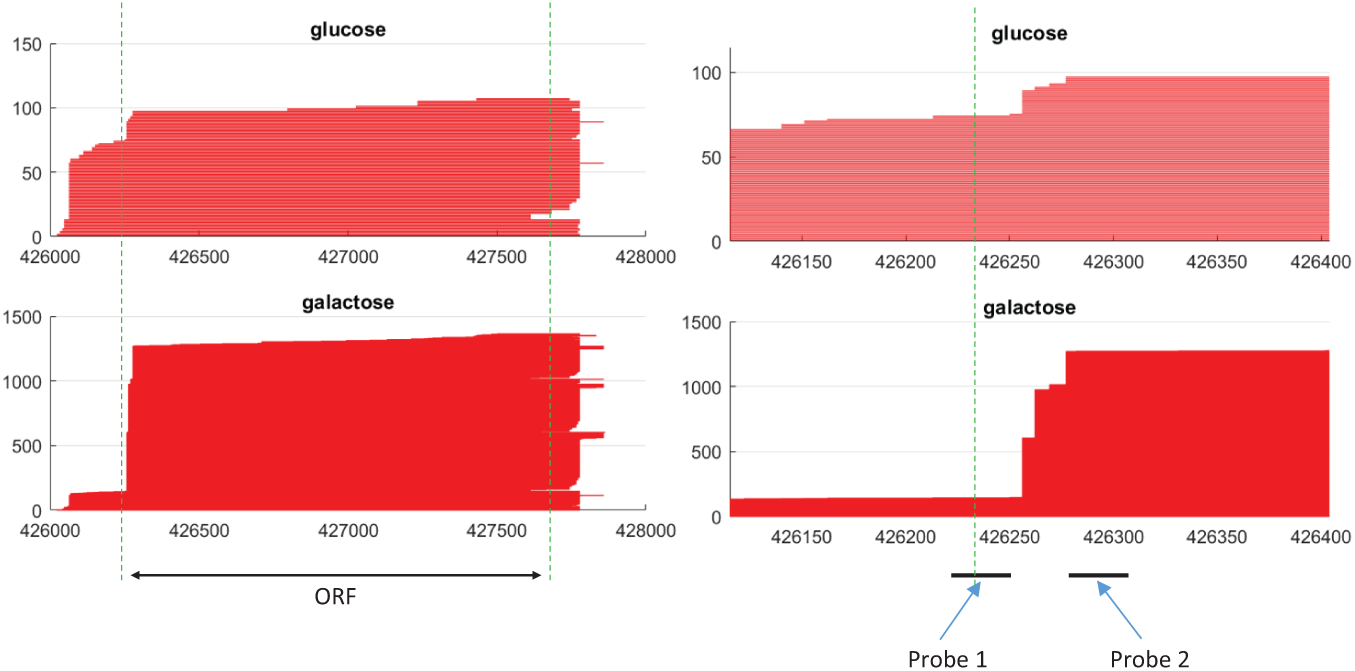
Determination of probe location for mRNA isoform profiling. mRNA isoform data for a yeast gene RGL1 (YPL066W) in glucose (top row) and galactose (bottom row) are shown at different zoom levels (zoom-out view on the left column and zoom-in view on the right). The x-axis represents the genomic coordinates around RGL1 on Chromosome XIV. Green vertical lines mark the ORF boundary. mRNA isoforms published in Pelechano et al. (38) are represented by red horizontal lines stacked vertically in the order of start coordinate. As shown, the transcriptional profile of RGL1 changes dramatically from glucose (top row) to galactose (bottom row). The target locations of Probe 1 and Probe 2 are shown as black bars.

**Supplementary Table S1.**
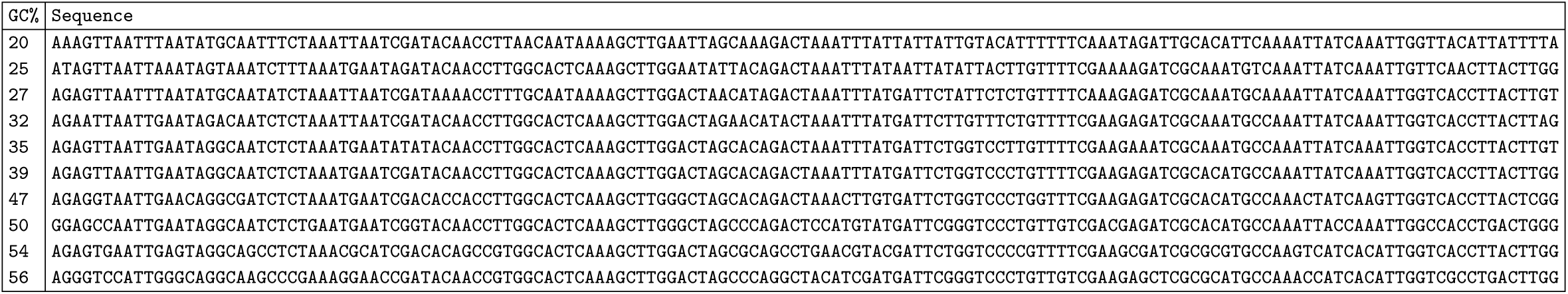
DNA sequence in the region of nucleosome -2.

**Supplementary Table S2.**
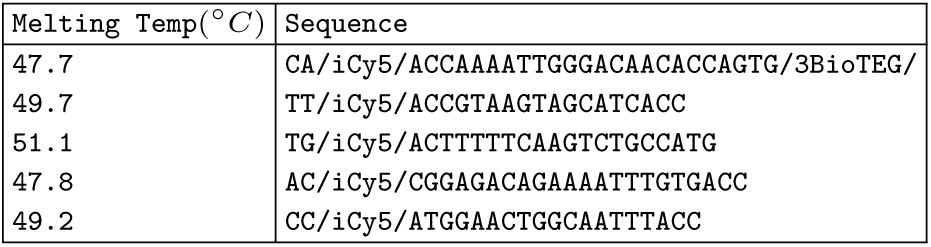
DNA sequence for yEVenus probes of similar melting temperature. Probes were designed to have similar GC content and melting temperature. The first probe is used for all of the single probe experiments.

**Supplementary Table S3.**
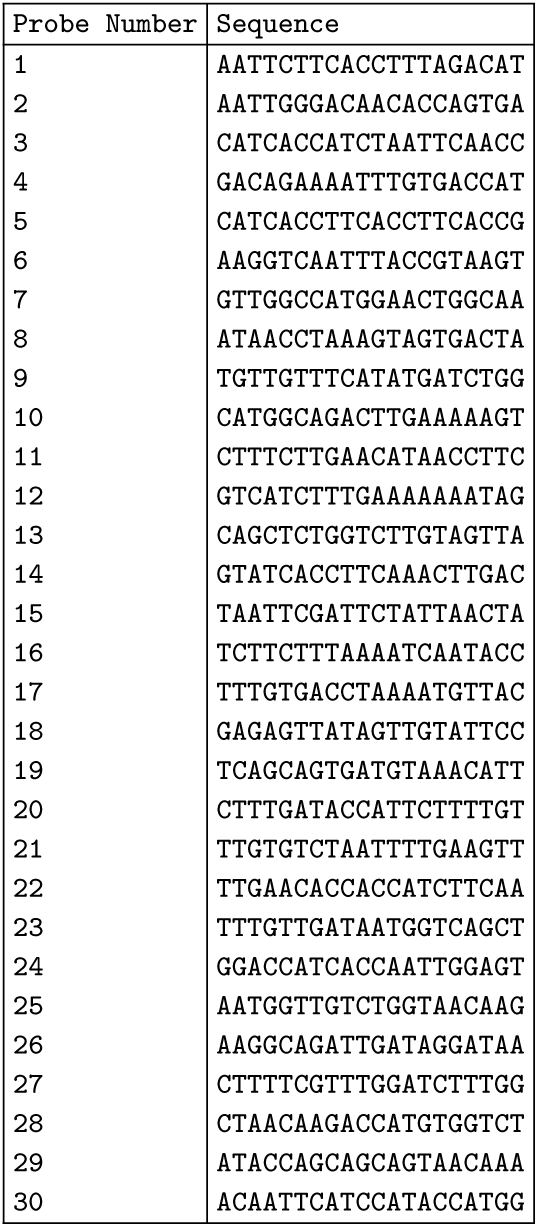
Probe sequences. Listed are sequences of probes used against yEVenus mRNA sequence(32). These sequences are identical to all but two used in a previous study(68).

**Supplementary Table S4.**
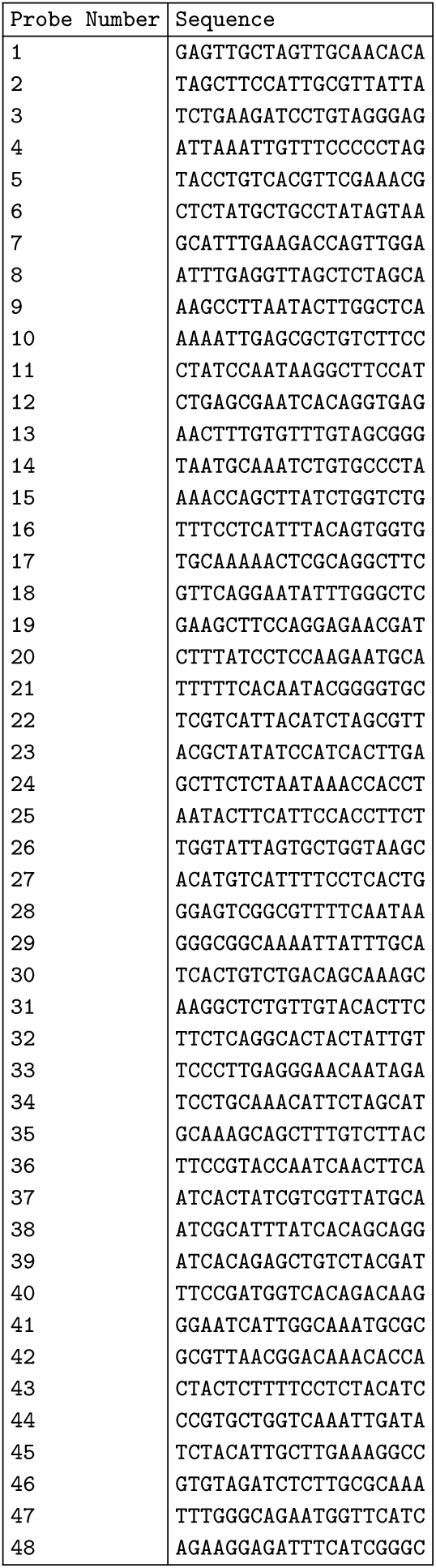
Probe sequences. Listed are sequences of probes used against KAP104 mRNA sequence. These sequences are identical to those used in a previous study(36).

**Supplementary Table S5.**
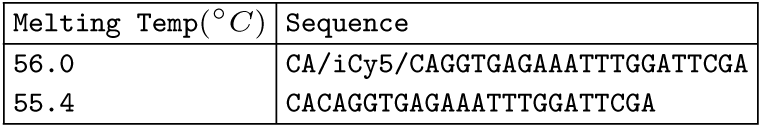
DNA sequence for KAP104 probes used in sFISH experiments.

**Supplementary Table S6.**
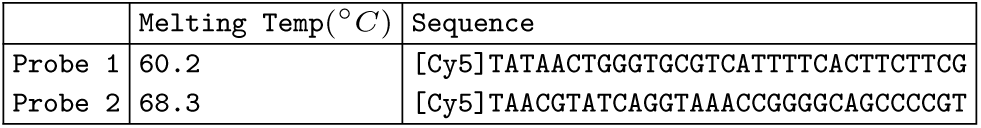
DNA sequence for RGL1 probes.

